# Climate change and socioeconomic vulnerability: The Carpathian Basin as a potential hotspot in the dissemination of *Dirofilaria repens* in Europe

**DOI:** 10.64898/2026.06.30.735485

**Authors:** Ágnes Csivincsik, Eszter Nagy, Izabella Zám, Tamás Tari, István Kucsera, Gábor Nagy, Tamás Sréter

**Affiliations:** One Health Working Group, Kaposvár Campus, Hungarian University of Agriculture and Life Sciences, 36 Guba Sándor Str., 7400, Kaposvár, Hungary; Institute of Wildlife Biology and Management, University of Sopron, 9400, 4 Bajcsy-Zsilinszky Str., Sopron, Hungary; National Reference Laboratory of Medical Parasites, National Centre for Public Health and Pharmacy, 1097, 2-6 Albert Flórián Str., Budapest, Hungary

**Keywords:** *Dirofilaria repens*, Carpathian Basin, Danube corridor, MaxEnt, socioeconomic factor, stray dogs, spatial Empirical Bayes smoothing, Bivariate Local Indicators of Spatial Association

## Abstract

**Background:** *Dirofilaria repens* is a zoonotic parasite expanding unnoticed across Europe due to climate change. We hypothesised that in this process, the Carpathian Basin has a facilitating effect.

**Methods:** Using 426 georeferenced European cases, the probability of infection occurrence was determined in relation to climatic factors, surface water availability, regional social deprivation, and stray dog population density. To analyse the potential impacts of ecological and social factors (deprivation and stray dog population density), the MaxEnt algorithm, and spatial Empirical Bayes smoothing and Bivariate Local Indicators of Spatial Association (BiLISA) index calculation were employed, respectively.

**Results:** MaxEnt analysis revealed that the mean warmest month temperature (22.8 - 25.1 °C), winter mean minimum temperature (> -2.1 °C), and summer precipitation (28.6 - 231 mm) have the strongest impact on the probability of the parasite’s occurrence in Europe. Social factors have significance in the eastern Balkans and the Carpathian Basin, but not in Western Europe. The Carpathian Basin appears to be a hotspot, similar to Mediterranean coastal areas. Furthermore, the Danube Valley acts as an ecological corridor for subtropical vector-borne parasites.

**Conclusions:** Our findings confirm that summer warmth is the primary ecological driver of the parasite’s range expansion, which is facilitated by the Carpathian Basin due to climatic and socioeconomic conditions.

## Background

Human dirofilariasis, caused by *Dirofilaria repens*, is a rapidly expanding parazitozoonosis in Europe [1]. It is agreed that the coexistence of canid reservoirs, vector populations, and appropriate conditions for larval development in vectors is essential for the formation of endemicity [2].

As a vector-borne infection transmitted by mosquitoes, *D. repens* is highly sensitive to climate change. In Europe, *Aedes vexans*, *Aedes sticticus*, *Aedes (Ochlerotatus) caspius*, *Culex pipiens* s.l., *Anopheles maculipennis* s.l., and *Anopheles hyrcanus* are the most frequently confirmed to be the vectors of the parasite [1]. The predominant vector species, *Ae. vexans* is a common floodwater mosquito [3,4]. Its primary biotopes are temporary freshwater basins created by seasonal river floods, snowmelt, or heavy rains. Females lay eggs on damp soil near these ephemeral ponds, which can remain viable for years until inundation triggers massive larval hatching. *Aedes vexans* can tolerate drying, as its larvae have shown remarkable survival even in humid soil [5]. It is one of the most widespread pest mosquitoes globally. Due to the wide tolerance to humidity, *Ae. vexans* occupies wet-prairie habitats in North America [6] and predicted to emerge in North African territories [7]. Owing to its migration potential, freshly hatched *Ae. vexans* individuals can fly up to 15 km from their breeding sites seeking blood meals [3]. Together with *Ae. caspius*, which is also a floodwater mosquito, *Ae. vexans* benefits from stormwaters of extreme weather events and artificial flooding, e.g. ricefields [4].

*Culex pipiens* s.l. is widely distributed in temperate regions across the northern hemisphere, [8,9], where it usually breeds in anthropised habitats. In Europe, the *Culex pipiens* complex (*Cx*. *Pipiens* s.l.) consists of several species, including *Culex pipiens* s.s. (Linnaeus, 1758) and *Culex quinquefasciatus* (Say, 1823) [10]. In addition, for *Cx*. *pipiens* s.s., two biotypes are recognized *Cx*. *pipiens* biotype *pipiens* and *Cx*. *pipiens* biotype *molestus*. The *Cx*. *pipiens* biotype *pipiens* is typically associated with aboveground habitats, such as standing water in clogged gutters or artificial containers, and may exhibit a bird-biased blood-feeding pattern [11,12] (Kim et al., 2018; Vinogradova, 2000). In contrast, *Cx*. *pipiens* biotype *molestus* (Forskål, 1775) is frequently found in underground habitats, such as basements, tunnels, and sewers. These two *Cx*. *pipiens* biotypes are capable of interbreeding [9–11].

Among the potential vectors of *D. repens*, *An. maculipennis* s.l. depends on the availability of open surface waters, woodlands and shrublands, and farm buildings for a successful life cycle [13]. Although climatic conditions are suitable for its maintenance on most of the European continent [14], the mosquito’s land use preference results in fragmented distribution [15]). Current climate change facilitates the northward distribution of *Anopheles* mosquitoes [15,16]; conversely, within their historical Mediterranean range, their occurrence probabilities may decline [14].

Some studies have raised the possibility that the epidemiological dynamics of *D. repens* might be affected by the socioeconomic deprivation of the local community, primarily due to the surrounding dense stray dog population [17,18]. Besides free-roaming dogs, marginalised social groups face other risk factors, such as poor education, low public health standards, and financial vulnerability, all of which exclude even owned dogs from protection against microfilarial infection [19,20]. Regarding *Dirofilaria immitis*, the socioeconomic factor is well-documented [21–23], while it has not been evaluated directly in the case of *D. repens*.

The Carpathian Basin is located in the centre of the European continent, geographically bounded by the Carpathian Mountains, the Alps, the Dinaric, and the Balkan mountains [24,25]. This area has played a central role in human migration [26,27]. Biogeographical studies have proven that both plant [28,29] and animal species [30] utilised the Carpathian Basin as a dispersion route from the Mediterranean zone northward. Furthermore, this central zone of Europe has also contributed to the spread of diseases [31–33].

Throughout human history, the Carpathian Basin has been a melting pot of nations at the southeastern entrance to Europe. Therefore, its disease-accumulating and transmitting roles were a direct consequence of its geographical location. Currently, another aspect of the Carpathian Basin’s uniqueness is highlighted. Global climate change affects certain regions differently. Although there is a consensus that the Arctic and mountainous regions are the most affected [34], the Carpathian Basin is also considered an extremely vulnerable region of Europe [25]. According to predictions, severe warming and aridification are expected [35,36], alongside a parallel increase in flood risk, due to the projected elevation of precipitation in the Carpathians [25,37]. These expected climate alterations might facilitate the reproduction of disease vectors, predominantly the floodwater mosquitoes, like *Ae. vexans* and *Ae. caspius*, as has been observed in flood-impacted regions of Europe [3,4].

Besides the climatic effect, the epidemiological risk of certain diseases is enhanced by the socioeconomic vulnerability of marginalised societal strata living in the region. Before the election on 12 April 2026, the autocratic Orbán regime ruled Hungary for 16 years, resulting in the social deprivation of the lowest-income communities and the widening of the poverty gap [38]. This long-standing kleptocracy caused devastating damage to both welfare redistribution [38,39] and the overall educational attainment of the population [40,41], leaving wide segments of the society behind, excluded from regular health service, quality education and the chance of social mobility [38,40]. Although mitigating the poverty gap is among the priorities of the new government, the healing of Hungarian society is expected to be a long process before this socioeconomic health risk is successfully managed.

Our study aimed to determine the risk factors of human *D. repens* infection in Europe, with a particular focus on Hungary, to highlight the potential role of the Carpathian Basin in the dissemination of the parasite.

## Methods

To achieve the objectives of our research, a sequential data analysis process was conducted, as demonstrated in the flowchart in Figure 1. The baseline dataset, utilised in this study, and the supplementary files are available in the Zenodo repository (https://zenodo.org/records/21054890).

**Figure 1.**
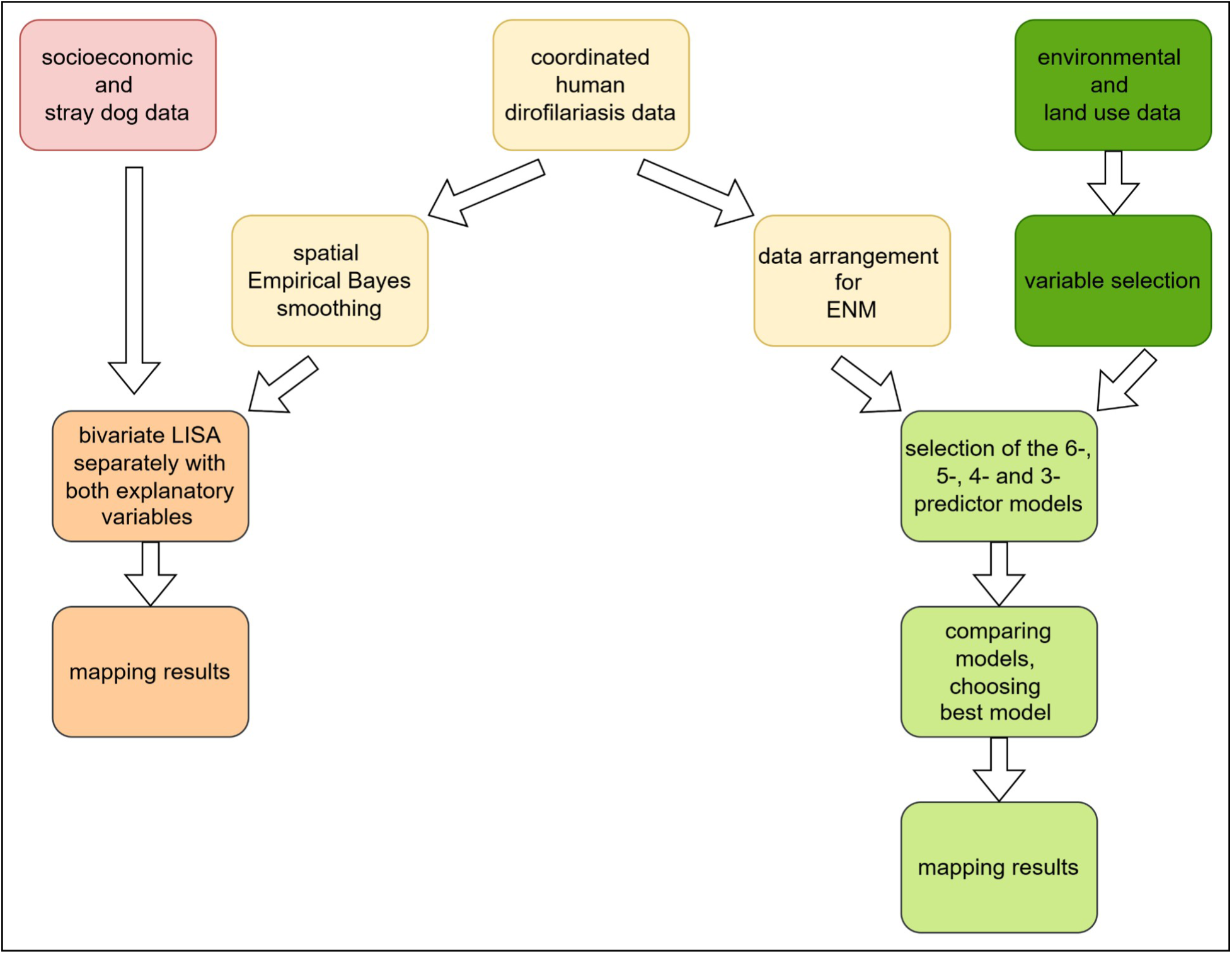
Methodological flowchart showing the data analysis processes

### Ecological Niche Modelling of HD

MaxEnt software is a multi-purpose machine learning tool [42] that models the geographic distribution of a species using georeferenced occurrence points. The method can also be used with high confidence for disease mapping [43–45]. The occurrence points, along with the environmental variables, serve as the input to the ENM. By selecting environmental variables, a prediction can be made that indicates the suitability of habitats for the occurrence of a given species or disease with high accuracy [46,42].

#### Occurrence data

For the modelling, we used human cases of dirofilariasis (*Dirofilaria repens*) (n = 1199) that were documented in Europe between 1885 and 2025 [1]. A significant proportion of the recorded cases occurred between 1990 and 2025 (n = 1013). In the case of Hungary, we used retrospective data obtained from the database of the National Reference Laboratory of Medical Parasitology (NLR, Department of Bacteriology, Mycology and Parasitology, National Centre for Public Health and Pharmacy). The Hungarian database contains epidemiological information on all *D. repens* specimens that were submitted to the NLR for parasitological laboratory confirmation of suspected cases. The NLR used morphological features [47,48] to identify *D. repens*. For this study, only aggregated epidemiological data, numbers of patients by postal codes, were assigned to the authors. Since all these data have been utilised in the regular communication activities of the National Centre for Public Health and Pharmacy and no identifiable personal patient information was disclosed, this study did not require specific ethical approval or special permission. Before processing, all double items (Hungarian NLR data published by Hattendorf & Lühken [1]) were removed from the dataset.

To model disease occurrence, the ENM method was used. For settlement-level data, we used the settlement’s central coordinate in the analyses. In the first step, spatial thinning (minimum distance of 5 km) was applied to the entire set of points to reduce spatial autocorrelation. The study area (M-space) was defined as a 100 km buffer zone around the retained points. This constituted the environmental space in which the discourse could thereby reduce bias from background sampling during the learning of the applied algorithm.

#### Environmental variables

Twelve bioclimate variables and one land-cover variable were used for the modelling. The climate data, covering the period 1992-2020, were obtained from the ClimatEU database (https://sites.ualberta.ca/~ahamann/data/climateeu.html, accessed on 14 June 2026). The only land-cover variable was the proportion of surface water, determined using the CORINE Land Cover 2018 (CLC2018) database (Table 1).

**Table 1.**
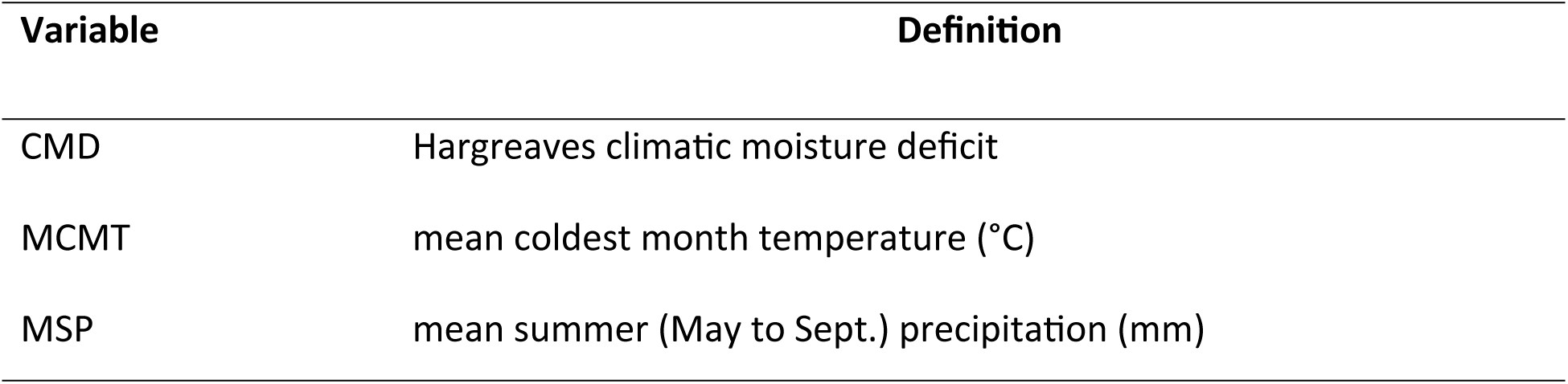
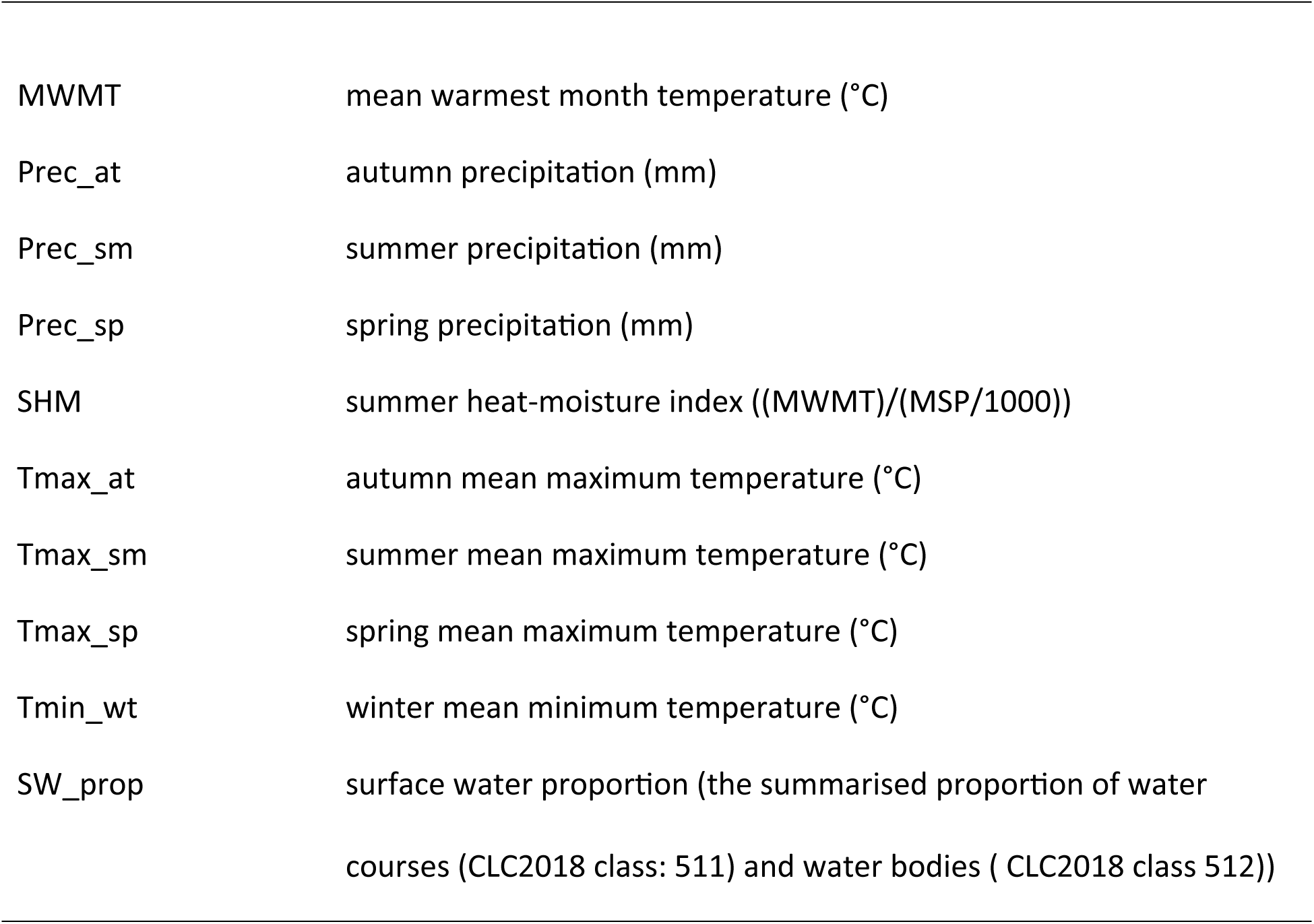
Potential explanatory variables involved in the Ecological Niche Modelling of human dirofilariasis in Europe.

| Variable | Definition |
| --- | --- |
| CMD | Hargreaves climatic moisture deficit |
| MCMT | mean coldest month temperature (°C) |
| MSP | mean summer (May to Sept.) precipitation (mm) |
| MWMT | mean warmest month temperature (°C) |
| Prec_at | autumn precipitation (mm) |
| Prec_sm | summer precipitation (mm) |
| Prec_sp | spring precipitation (mm) |
| SHM | summer heat-moisture index ((MWMT)/(MSP/1000)) |
| Tmax_at | autumn mean maximum temperature (°C) |
| Tmax_sm | summer mean maximum temperature (°C) |
| Tmax_sp | spring mean maximum temperature (°C) |
| Tmin_wt | winter mean minimum temperature (°C) |
| SW_prop | surface water proportion (the summarised proportion of water<br>courses (CLC2018 class: 511) and water bodies ( CLC2018 class 512)) |

The data were used as raster projections. Since the climatic and land cover variables were available at different spatial resolutions, it was necessary to standardise the rasters to ensure modelling accuracy and compatibility across layers. For this unified spatial analysis base, we chose a resolution of ∼2km resolution (75 arcsec).

The accuracy of the model can be greatly affected by multicollinearity among variables [49]. We have addressed this issue with two-stage filtration, based on the pairwise Spearman correlation coefficients (threshold: |ρ|<0.8), then using the variance-inflation factor (VIF) (threshold: VIF< 5) [50]. As a result of the filtering, six variables remained for modelling: mean warmest month temperature(MWMT), winter mean minimum temperature (Tmin_wt), summer heat-moisture index (SHM), amount of summer precipitation (Prec_sm), spring precipitation (Prec_sp), and proportion of surface water (SW_prop). Collinearity screening was performed with the usdm [51] and terra [52] R packages.

#### Model building and model selection

The Ecological Niche Model (ENM) building was carried out using the MaxEnt algorithm [42], which estimates the relative probability of species occurrence based on available environmental conditions, using the principle of maximum entropy. To optimise and evaluate the model, we used the ENMeval R package [53], which systematically tests combinations of feature class (FC) and regularisation factor (RM) and evaluates the model’s predictive performance using spatial partitioning methods.

The model selection was carried out using a cascading variable reduction strategy. Four nested sets of variables (6-, 5-, 4-, and 3-predictor models) were evaluated, with stepwise removal of the least informative variables. The order in which the variables were removed was determined based on three combined criteria: per cent contribution (PC), permutation importance (PI) and jackknife method. A variable was removed when all three indicators showed low values simultaneously [54].

A total of 120 model configurations were tested using the same settings, such as six feature-class combinations (L: linear, Q: quadratic, H: hinge, LQ: linear-quadratic, LH: linear-hinge, LQH: linear-quadratic-hinge) and five regularisation factors (RM = 1–5). During the process, we used checkerboard spatial partitioning with a 4 × 4 grid cell aggregation. This method divided the study area into regular, complementary blocks, which ensured the spatial independence of the training and validation data. The number of background points was 50,000. Model building was performed using MaxEnt software (version 3.4.4) within the ENMeval framework, with the terra and rJava R packages [55].

#### Model evaluation and comparison

The best model selection was based on the AICc (corrected Akaike Information Criterion), which accounts for both the model’s fit and complexity. A lower AICc indicates a better model. If the delta AUC was < 2 between two models, they were considered statistically equivalent [56]. The following indicators were used to evaluate the models:

- AUC (Area Under the ROC Curve): measures the model’s ability to discriminate, i.e. its ability to distinguish between suitable and unsuitable areas. A model was considered a failure if 0.5<AUC<0.6, poor if 0.6<AUC<0.7, fair if 0.7<AUC<0.8, good if 0.8<AUC<0.9, or excellent if 0.9<AUC [57].
- CBI (Continuous Boyce Index): evaluates the calibration of the model, i.e. the extent to which the predicted fitness values correlate with the density of the points of occurrence. A CBI > 0.7 is acceptable; a CBI> 0.9 indicates excellent calibration [58].
- 10% omission rate: the proportion of known points of occurrence that the model mistakenly indicates as unfit at the 10th percentile threshold. The ideal value is approximately 0.10 [59].
- Number of coefficients (ncoef): the number of parameters in the model, which is a direct indicator of the model complexity. A lower coefficient indicates a lower risk of overfitting [60].
- w.AIC (Akaike weight): the relative probability of a given model configuration among all tested configurations. A value> 0.9 indicates a clear selection [56] (Burnham and Anderson, 2002).

For a final comparison between the variable sets, the selected best configurations were also run with 10x cross-validation in MaxEnt. During the run, the sampling points were weighted 75:25 between the M-area (calibration area) and the non-M-area (projection area) to focus model training on the biologically relevant background [61]. From the results obtained, additional indices were formed for the selection of the final model:

- Test AUC: the mean AUC calculated on test sets of 10x cross-validation.
- TSS (True Skill Statistic): the sum of sensitivity + specificity-1, calculated on the threshold of maximum training sensitivity + specificity. TSS > 0.6 indicates good predictive performance [62,63].

#### Criteria for choosing the final model

The final model was selected by weighing the discriminating power (AUC, TSS), calibration (CBI), omission rate, and model complexity (number of coefficients). Since the aim of the study was a climate-based characterisation of the current prevalence of human dirofilariasis, we preferred discriminatory performance over transferability.

#### Types of response curves

The marginal response curves of the final model were interpreted within the framework of ecological niche theory [64,65]. Unimodal (with optimum) response curves are expected for direct environmental variables, and monotonic (saturation-like) response curves are expected for limiting factors [66]. The thresholds of the response curves of the variables in the final model were determined from the results in the MaxEnt lambda files.

### Socioeconomic analysis

We examined the disease and its socioeconomic background at the second level of the Nomenclature of Territorial Units for Statistics (NUTS 2), which is used as a reference in Europe. The analysis included the following 28 European countries: Austria, Belgium, Bulgaria, the Czech Republic, Denmark, North Macedonia, Finland, France, Greece, the Netherlands, Croatia, Ireland, Lithuania, Hungary, Montenegro, Germany, Norway, Italy, Poland, Portugal, Romania, Spain, Sweden, Switzerland, Serbia, Slovakia, Slovenia, and Türkiye.

For the analysis, we used only cases for which the location was specified with at least NUTS 2-level precision. Using the number of cases in each region and the region’s population, we estimated the relative risk of HD (HDSEBRR) using spatial Empirical Bayes smoothing. Since the population sizes of individual regions were highly heterogeneous, the raw rates were not suitable for a realistic analysis. Therefore, this approach allows us to correct for the resulting heterogeneity and variance in human population size. The basic principle of the method is to “scale down” the rates observed in each territorial unit toward a local reference value, calculated from a window defined by the spatial context of the given unit (e.g., neighbouring areas) [67].

The local reference average (μi) in the vicinity of the *i*-th spatial unit can be expressed as follows:

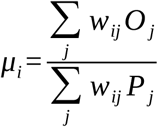

where *O_j_* is the number of events, *P_j_* is the at-risk population, and *w_ij_* are the spatial weights (binary neighbourhood matrix), where w_ij_ = 1 means that the given unit is itself part of the local window.

The local estimate of the prior variance is also based on data from the surrounding areas:

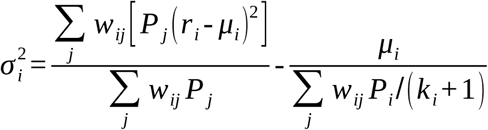

where *r_i_* is the observed proportion and *k* is the total population associated with the local window.

We have expressed the socio-economic characteristics of the regions using the rate of severe material and social deprivation as measured in Europe. This index shows the extent to which people lack the goods necessary and desirable for a decent standard of living. The index is expressed as a percentage, the socially deprived proportion of the population. Notably, this value is calculated for each country; therefore, it aligns with that country’s average standard of living. It is determined at the household and individual levels using various indicators (Table 2). We obtained population data for each region, as well as data on severe material and social deprivation, from the European Union’s Eurostat database (https://ec.europa.eu/eurostat/, accessed on 14 June 2026).

**Table 2.**
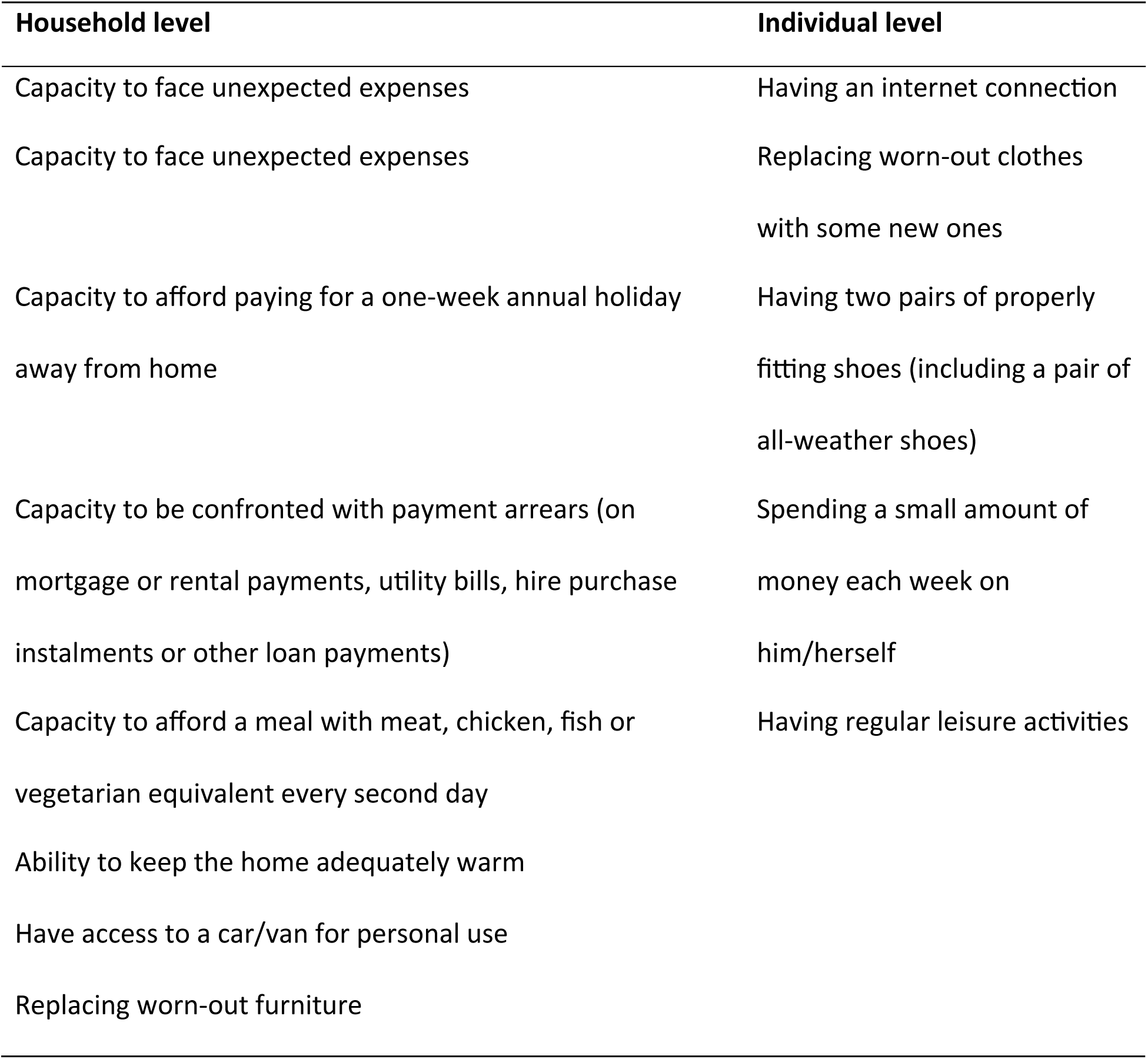
Indicators used to calculate the proportion of severe material and social deprivation in Europe.

| Household level | Individual level |
| --- | --- |
| Capacity to face unexpected expenses | Having an internet connection |
| Capacity to face unexpected expenses | Replacing worn-out clothes<br>with some new ones |
| Capacity to afford paying for a one-week annual holiday<br>away from home | Having two pairs of properly<br>fitting shoes (including a pair of<br>all-weather shoes) |
| Capacity to be confronted with payment arrears (on<br>mortgage or rental payments, utility bills, hire purchase<br>instalments or other loan payments) | Spending a small amount of<br>money each week on<br>him/herself |
| Capacity to afford a meal with meat, chicken, fish or<br>vegetarian equivalent every second day | Having regular leisure activities |
| Ability to keep the home adequately warm |  |
| Have access to a car/van for personal use |  |
| Replacing worn-out furniture |  |

To analyse the spatial relationship between the two variables, the Bivariate Local Indicators of Spatial Association (BiLISA) index was calculated. A bivariate spatial autocorrelation index indicates the extent to which two variables tend to cluster spatially. The bivariate LISA is a simple extension of the LISA function to two different variables, one related to location and the other to the average of neighbours [68]. The value of the index is positive if the two variables change together in space (High-High or Low-Low neighbourhood), and negative if they change in opposite directions (High-Low or Low-High neighbourhood). The significance level was evaluated by permutation testing with 10000 permutations. The following formula was used for the calculation:

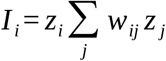

where *z_i_* is the standardised x value (explanatory variable) of the observation unit *i*, *w_ij_* is the element of the spatial weight matrix, and *z_j_* is the standardised y value (dependent variable) of the neighbouring unit *j*.

### Sray dogs and HD

There are no precise figures available on the number of stray dogs in Europe. As a result, we were only able to use estimated data for our analysis, which we compiled from the following website: https://www.esdaw.eu/stray-animals-by-country.html (accessed on 14 May 2026). Because this approximation was available only at the national level, a spatial interpolation was applied for the NUTS 2-level analysis. We distributed the estimated number of stray dogs in each country according to the relative areas of the regions within each country. This approach assumes a uniform spatial distribution, which does not hold in reality. Due to the lack of systematic subregional data on stray dogs, we deemed this method the best available approximation. When interpreting the results, it is therefore important to note that the regional distribution of stray dogs may differ from the actual situation, and that the analysis is suitable for identifying large-scale spatial trends rather than for providing precise risk estimates for individual regions. In both cases, for data sorting and spatial analysis, we used GeoDa (version 1.22.0.14) and QGIS (version 3.32, Lima).

## Results

Between 1999 and 2025, a total of 148 human *D. repens* infections were detected in Hungary, of which three were not georeferenced; therefore, 145 cases were analysed (Figure 2).

**Figure 2.**
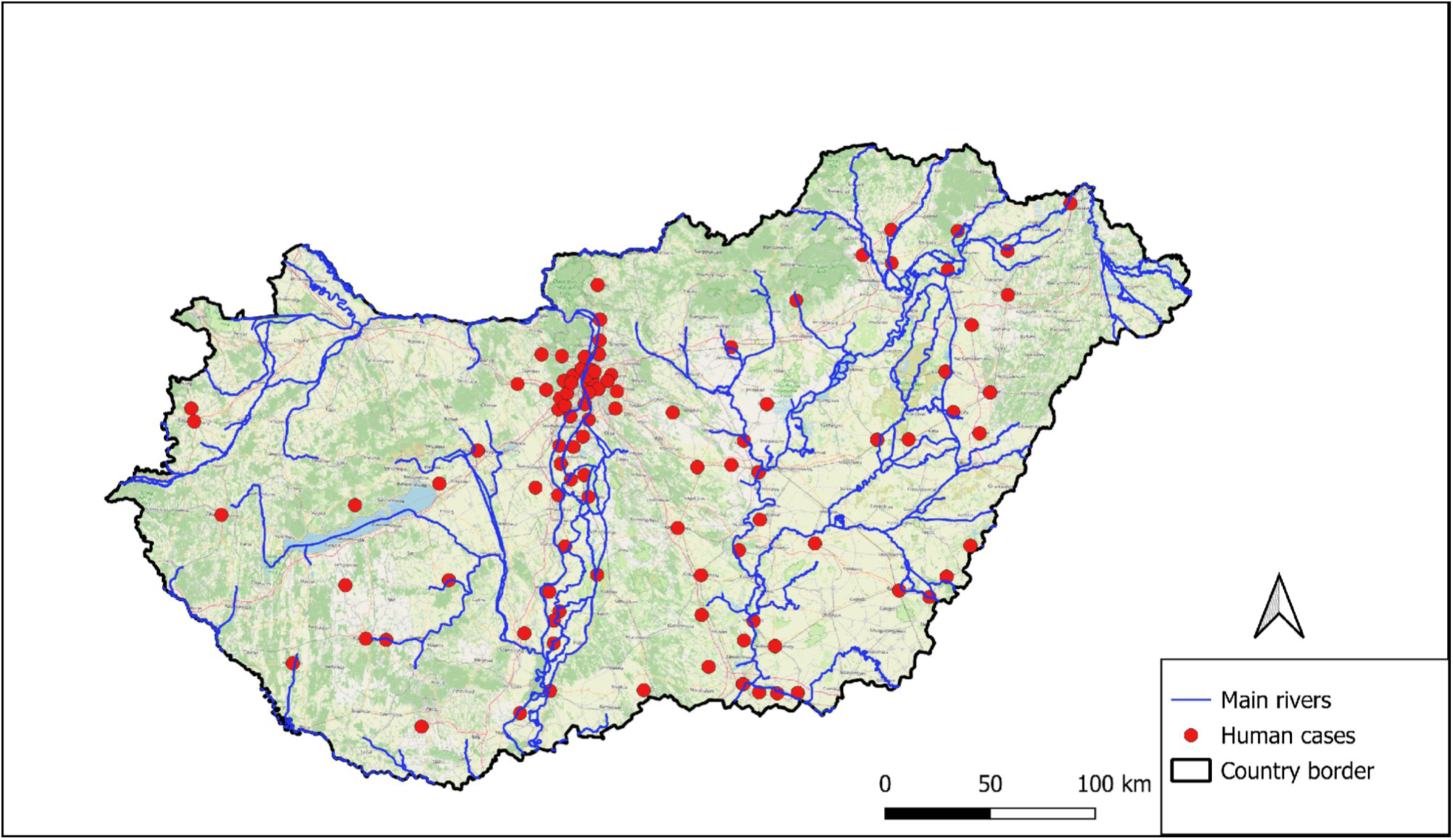
Locations of *Dirofilaria repens-*infected human cases detected between 1999 and 2025 in Hungary. (Note: The map presents 145 cases. The western and eastern parts of the country belong to the Danube and the Tisza River catchment, respectively.)

### Ecological Niche Modelling

To explore the probability of HD, 1119 records were used in ecological niche modelling, of which 145 were from Hungary. The cases are concentrated primarily in Central and Southern Europe, and within that region, most notably in the area between 8 and 21 degrees east longitude and 40 and 50 degrees north latitude. After excluding cases outside the used layers, applying thinning, and trimming duplicates, 426 cases remained. This point set was applied to estimate the probable occurrence of the disease in Europe (Figure 3).

**Figure 3.**
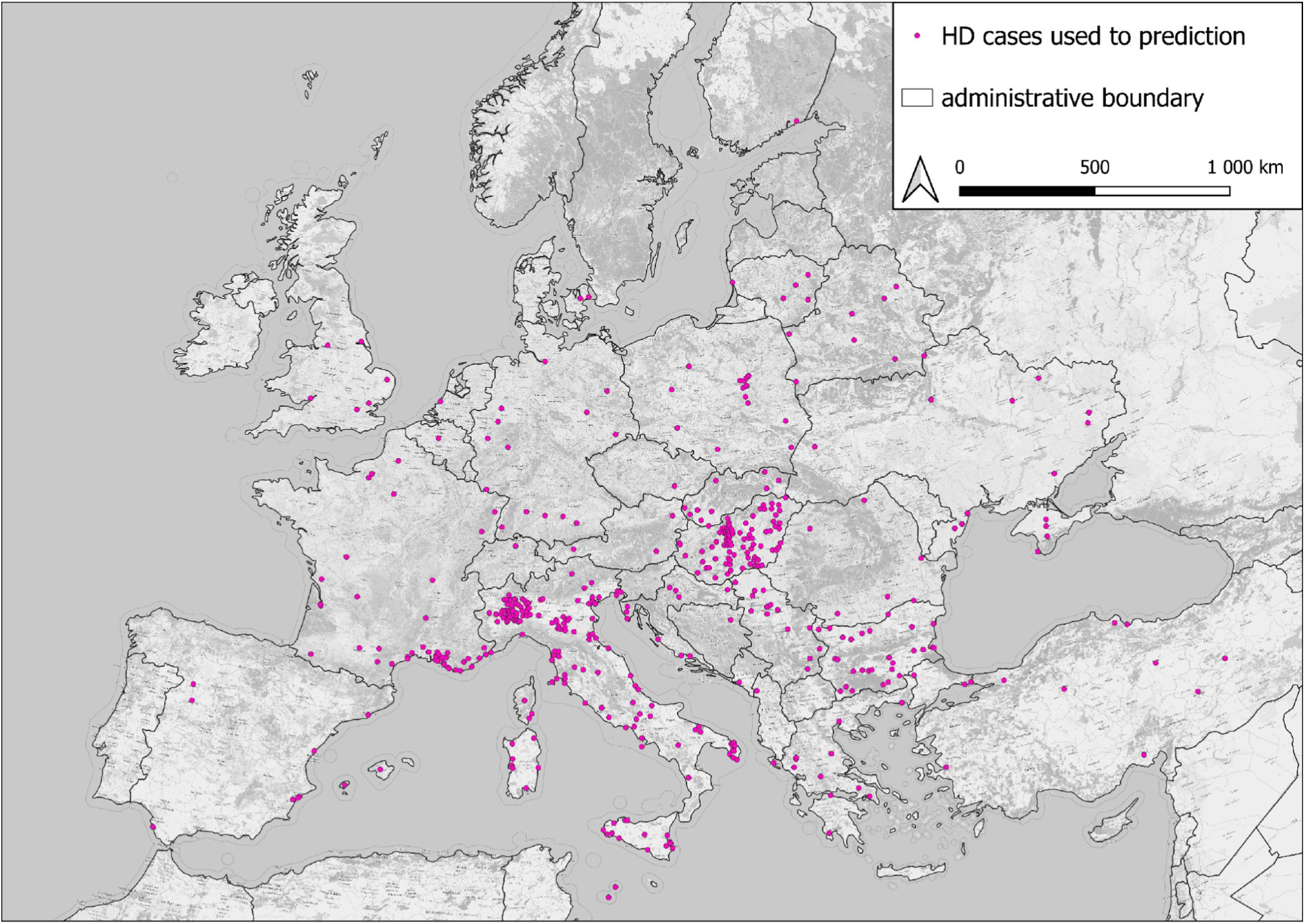
Geographical distribution of HD in Europe. (Note: The pink circles represent the points from another study [1] and from the Hungarian database of the National Centre for Public Health and Pharmacy.)

Of the initial 13 explanatory variables, seven were rejected due to collinearity. After selection, the following variables were used in the ENM: mean warmest month temperature (MWMT, VIF = 3.48), summer precipitation (Prec_sm, VIF = 2.72), spring precipitation (Prec_sp, VIF = 1.59), summer heat moisture index (SHM, VIF=1.49), surface water proportion (SW_prop, VIF = 1.02), and winter mean minimum temperature (Tmin_wt, VIF = 2.14).

A total of 120 models were examined during the analysis. The final model’s set of variables consisted of three predictors: MWMT, Tmin_wt, and Prec_sm. Since the aim of the study was to ecologically characterise the current distribution of human dirofilariasis rather than to model future scenarios, we prioritised discriminatory performance over transferability when selecting the best model. We stepwise excluded the least important explanatory variables from the models based on percentage contribution (PC), permutation importance (PI), and jackknife analysis.

From the initial model containing six explanatory variables, we first excluded SW_prop (PC = 1.0%, PI = 0.7%), which had the lowest jackknife loss among all variables. In the second step, we excluded Prec_sp (PC = 3.8%, PI = 3.2%) and subsequently SHM, since its percentage contribution in the four-variable model had dropped to practically zero (PC = 0.08%). Since its regularised training gain was 0.001, the SHM removal caused a negligible effect on the model. During the model selection process, the optimal configuration of the final selected model was a linear hinge (LH) object class with a regularisation coefficient of three.

This model performed the highest validation AUC (0.881), the lowest 10% omission rate, the highest TSS (0.627), and strong calibration (CBI = 0.936) among all variable configurations (Table 3).

**Table 3.** Results of model selection for *Dirofilaria repens.* (Note: The models were evaluated using checkerboard partitioning (aggregation factor: 4 × 4), with 50000 background points and 30 feature class/regularisation coefficient combinations per variable set.)

| indices | model1 | model2 | model3 | model4 |
| --- | --- | --- | --- | --- |
| number of variables | 6 | 5 | 4 | 3 |
| FC/RM | LQ/1 | LQ/1 | LH/3 | LH/3 |
| ncoef | 10 | 10 | 64 | 50 |
| AICc | 13253.7 | 13274.2 | 13303.4 | 13265.7 |
| AUC | 0.8791 | 0.8763 | 0.8802 | 0.8812 |
| AUC SD | 0.027 | 0.027 | 0.030 | 0.030 |
| CBI | 0.968 | 0.968 | 0.929 | 0.936 |
| 10% omission | 0.101 | 0.105 | 0.103 | 0.099 |

(Note: FC/RM= feature class / regularisation multiplier; ncoef=number of coefficient; AICc=Akaike Information Criterion, corrected; AUC= Area Under the Receiver Operating Characteristic Curve; AUC SD=AUC Standard Deviation; CBI=Continuous Boyce Index; 10% omission=10 percentile training presence omission rate; LQ=linear-quadratic, LH=linear-hinge)

The performance of the final model was also confirmed by an independent 10-fold cross-validation run (test AUC = 0.875; TSS = 0.627; 10% omission rate = 0.106). The three retained variables contributed considerably to the best model (Figure 4). MWMT showed the highest PC (46.6%) and PI value (54.5%). Tmin_wt and Prec_sm showed similar permutation importance, 29.4% and 23.9%, respectively. The percentage contribution value of Tmin_wt was 2.5 times higher (32.1%) than that of Prec_sm (13.4%).

**Figure 4.**
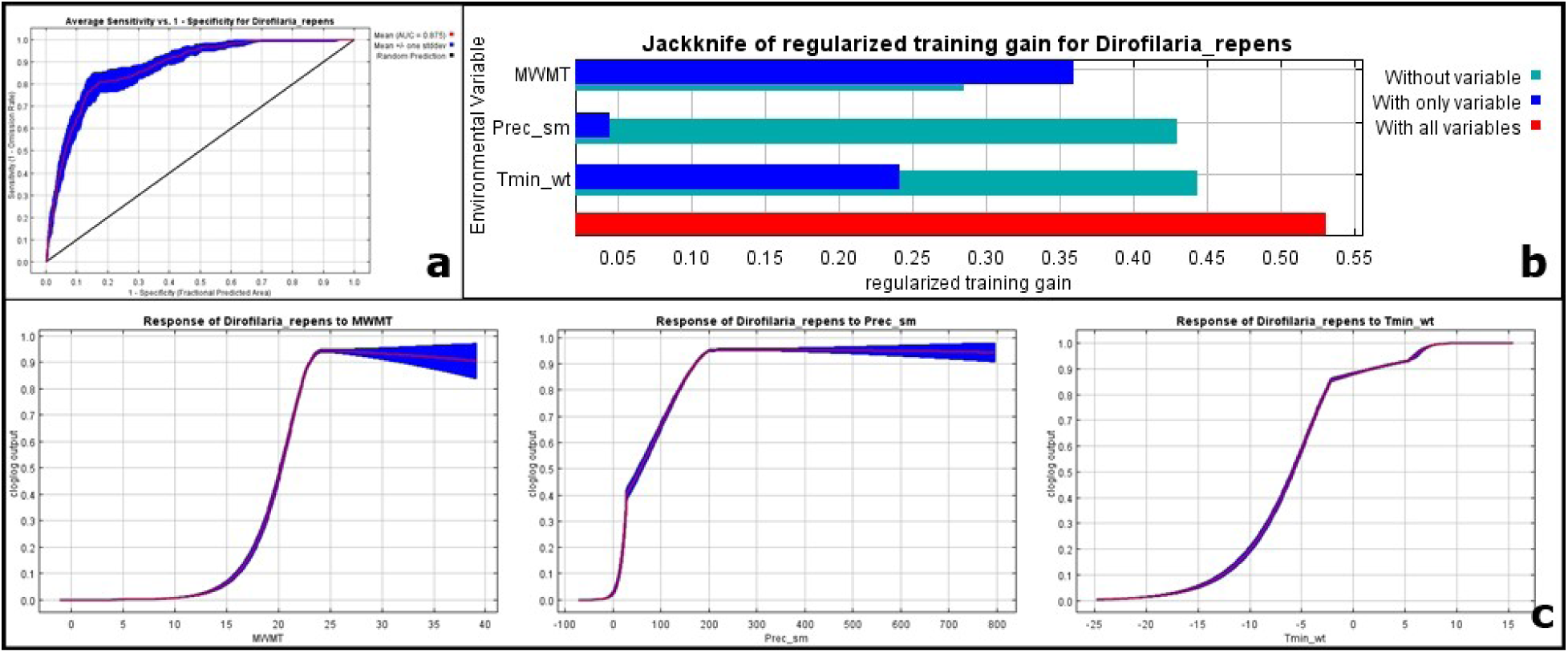
Results indicated the performance of the final model. (Note: The Receiver Operating Characteristic (ROC) curve and AUC value (**a**) in the investigated period (1911–2040). The jackknife test expresses the importance of different environmental variables (**b**). The response curves of the explanatory variables (**c**). The red lines express the average calculated from 10 model runs, while the blue shading indicates the standard deviation.)

The LH feature class identified sharp climatic thresholds for all variables, which proved consistent across the cross-validation folds (Figure 5). The unimodal marginal response curve of the mean temperature of the warmest month (MWMT) delineated an optimal zone between approximately 22.8 °C and 25.1 °C. Suitability decreases sharply below the lower limit, which could indicate the lower temperature limit for the development of *Dirofilaria repens* L3 larvae in the intermediate host. Above the optimum, suitability decreased moderately, suggesting that excessively high summer temperatures may limit transmission through reduced mosquito activity or survival. The Tmin_wt response curve exhibited a monotonic increase, with a sharp lower threshold at −2.1 °C but no upper threshold. The absence of an upper thermal limit indicates that, within the study area’s climatic range, winter temperature serves as a lower threshold (a limiting factor), so an optimum could not be determined. The summer precipitation (Prec_sm) had a unimodal response curve. This variable exhibited the model’s strongest single hinge effect. Below 28.6 mm, suitability dropped to nearly zero, suggesting a drastic reduction in the moist environment necessary for mosquito reproduction. Above the upper limit of the optimum (231 mm<), the suitability also decreases.

**Figure 5.**
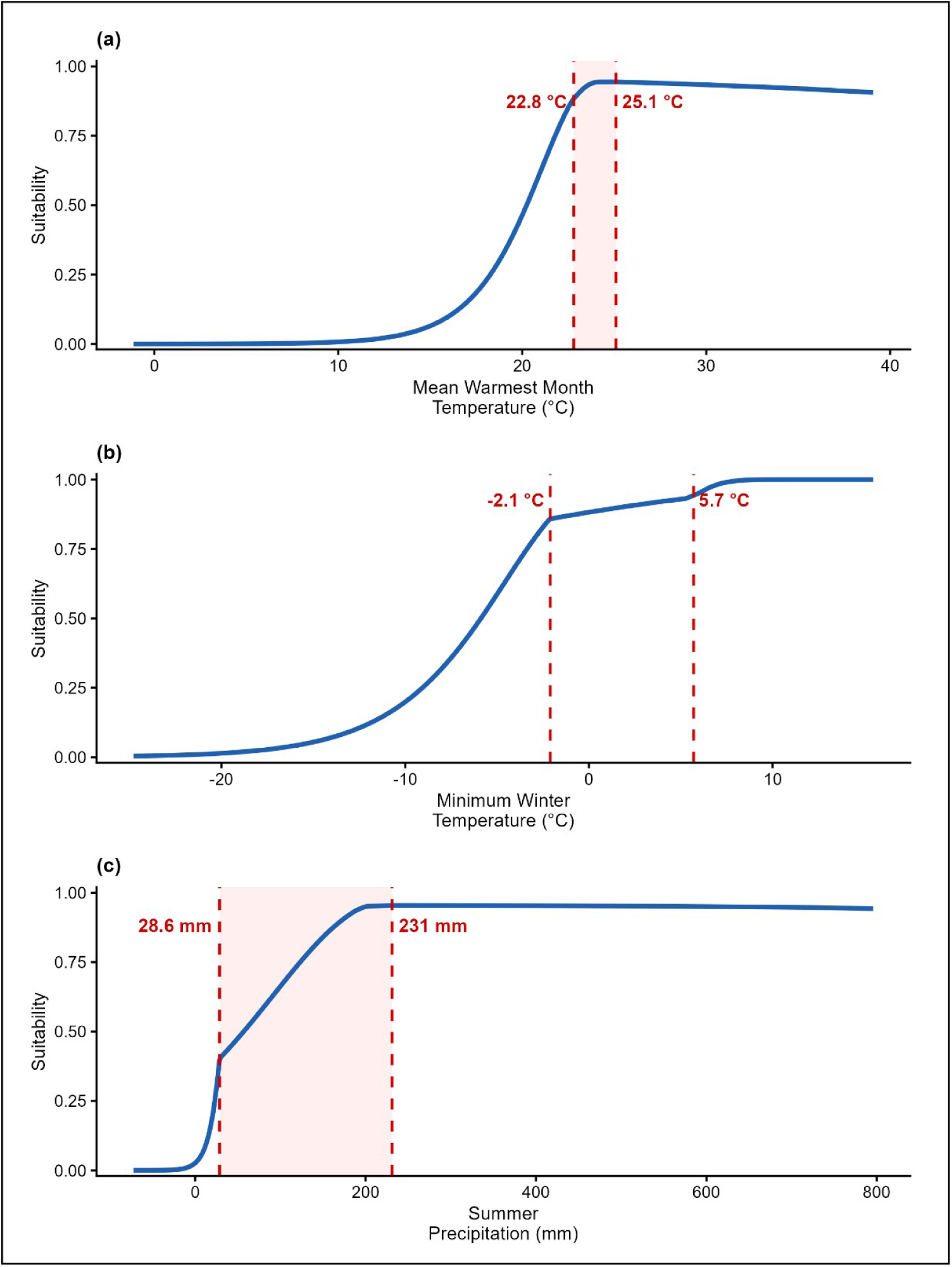
Marginal response curves of the final predictive model (LH, RM = 3) for *Dirofilaria repens*. (Note: Panel **(a)** shows the mean temperature of the warmest month (MWMT), panel **(b)** shows the winter minimum temperature (Tmin_wt), and panel **(c)** shows the summer precipitation (Prec_sm). The red dashed lines indicate the threshold values. In panels **(a)** and **(c)**, the red shaded area indicates the optimal zone between the two threshold values. The y-axis shows the predicted suitability value.)

The areas with the highest probability (0.7<) are concentrated along the Mediterranean coastline and around the Black Sea. Southern France, the Italian Peninsula (especially the Po Valley and southern regions), Greece, the coastal strip of the Balkans, the eastern coast of the Black Sea, and the complete territory of Hungary also show exceptionally high values (Figure 6).

**Figure 6.**
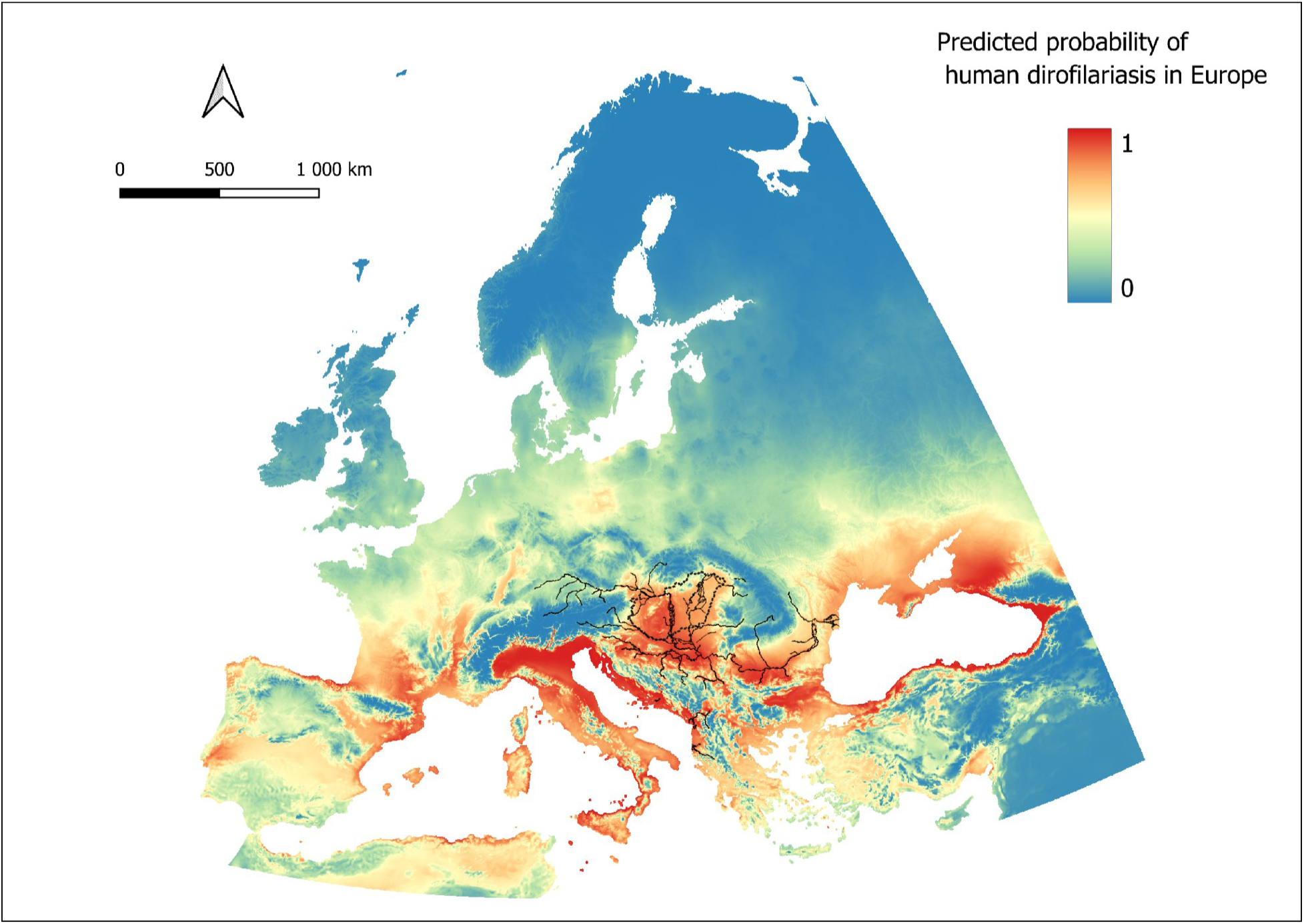
Prediction of human dirofilariasis in Europe and Hungary. (Note: The illustration is based on 426 cases with coordinates. The red colour indicates a higher likelihood of disease, while the blue area indicates a lower likelihood. The black dotted line denotes Hungary’s administrative border; black continuous line indicates Danube and its side stream tributaries.)

The medium-probability zone (0.3–0.7) forms a broad transitional zone in Western and Central Europe. This category covers a large part of the continent north of the Alps, as well as areas located in the immediate vicinity of the continent’s mountain ranges (e.g., the Dinaric Alps stretching across the central Balkans). In these areas, summer temperatures generally reach the lower threshold, but due to colder winter conditions, fewer parasite generations may develop in intermediate hosts. The pattern observed in the interior regions of the Iberian Peninsula is noteworthy, as predicted, suitability declines in some areas despite the otherwise high summer temperatures. This is consistent with the lower threshold of 28.6 mm for the Prec_sm variable: in the interior, semi-arid regions of the peninsula, summer precipitation does not provide sufficient breeding sites for mosquitoes, which may break the transmission.

The low-probability zones (<0.3) consisted of northern areas of Europe and the highest mountains. In these regions, both summer temperature and minimum winter temperature could limit high mosquito densities and thus the possible occurrence of human infections.

### Socioeconomic analysis

The bivariate LISA analysis revealed a significant spatial association between the spatial relative risk of human dirofilariasis (HDSEBRR) and material and social deprivation at the NUTS 2 level (Figure 7).

**Figure 7.**
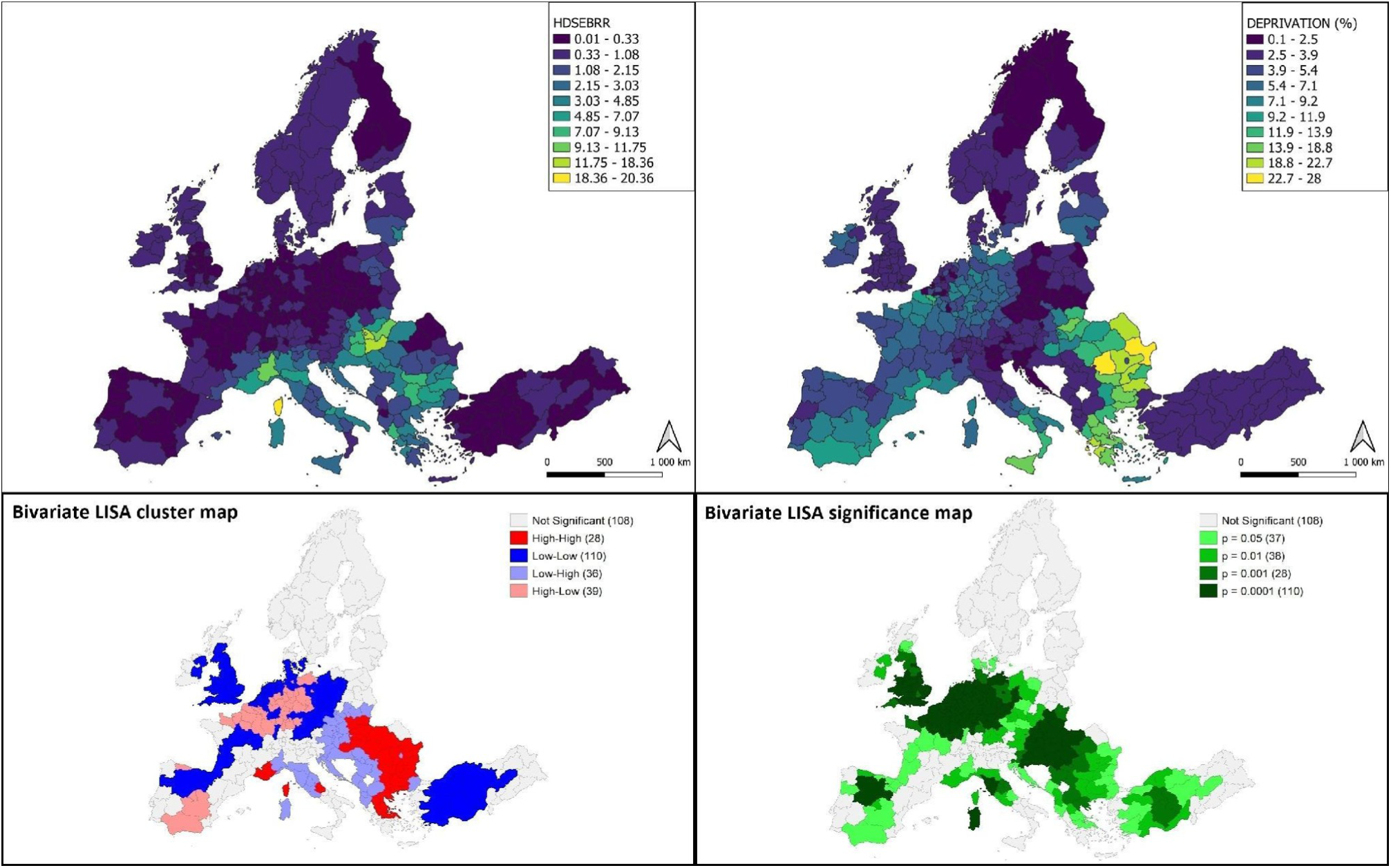
Spatial association between the spatial relative risk of human dirofilariasis and economic and social deprivation at the NUTS 2 level. (Note: Top left: HDSEBRR (spatial Empirical Bayes relative risk of human *Dirofilaria repens* infection). Top right: economic and social deprivation as a percentage of the population (DEPRIVATION). Bottom left: bivariate LISA cluster map (High-High: high deprivation and high infection risk; Low-Low: low deprivation and low infection risk; Low-High: low deprivation, high infection risk; High-Low: high deprivation, low infection risk; numbers of regions in parentheses). Bottom right: bivariate LISA significance map (number of regions per significance level in parentheses)).

The relative risk of HD (HDSEBRR) shows a clear southeast–northwest direction. Regions with the highest relative risk (7<) are concentrated in Southern Italy, Greece, the Balkans, and certain regions of Romania and Hungary. In contrast, most regions in Northern and Western Europe show low risk (1>). Economic and social deprivation follow a similar spatial pattern. The most deprived regions (>18%) are found in Central and Southeast Europe (Hungary, Romania, Bulgaria, Greece), while regions in Scandinavia, Germany, and the Benelux countries show values below 2.5%.

The LISA cluster map identified four cluster types in 213 of the 321 studied regions (66.4%). High-High clusters (28 regions), where high relative risk is coupled with high deprivation, are concentrated in Southeast Europe, particularly in the Balkans, Romania, and Greece, as well as most of Hungary. In contrast, the Low-Low clusters (110 regions), a combination of low risk and low deprivation, are found in Northern and Western Europe. The significance levels in these clusters are typically below p = 0.0001, indicating strong statistical support for the association.

The Low-High clusters (36 regions) are particularly noteworthy: they exhibit a low deprivation alongside high risk of dirofilariasis. They are primarily found in the northern and central regions of Italy, as well as in the western Balkans and the western and northern regions of the Carpathian Basin. This pattern suggests that in these economically more developed regions, the occurrence of the disease is determined not by social disadvantage but primarily by climatic suitability, in line with the results of the ENM analysis, which showed high suitability values for the Mediterranean and the Carpathian Basin.

The High-Low clusters (39 regions), high deprivation, low risk, are found mainly in the Atlantic climate zones of Western Europe and the Iberian Peninsula, where, despite relative social disadvantage, climatic conditions (low summer temperatures, extreme winter cold, or, conversely, extreme aridity) do not permit effective transmission.

### Number of stray dogs

The second bivariate LISA analysis examined the spatial relationship between the estimated relative risk of human dirofilariasis and the estimated number of stray dogs (Figure 8). The estimated number of stray dogs shows dramatic regional differences: the highest values (>150,000 per region) are found in Romania, while in most regions of Western and Northern Europe, the estimated number is below 5,000.

**Figure 8.**
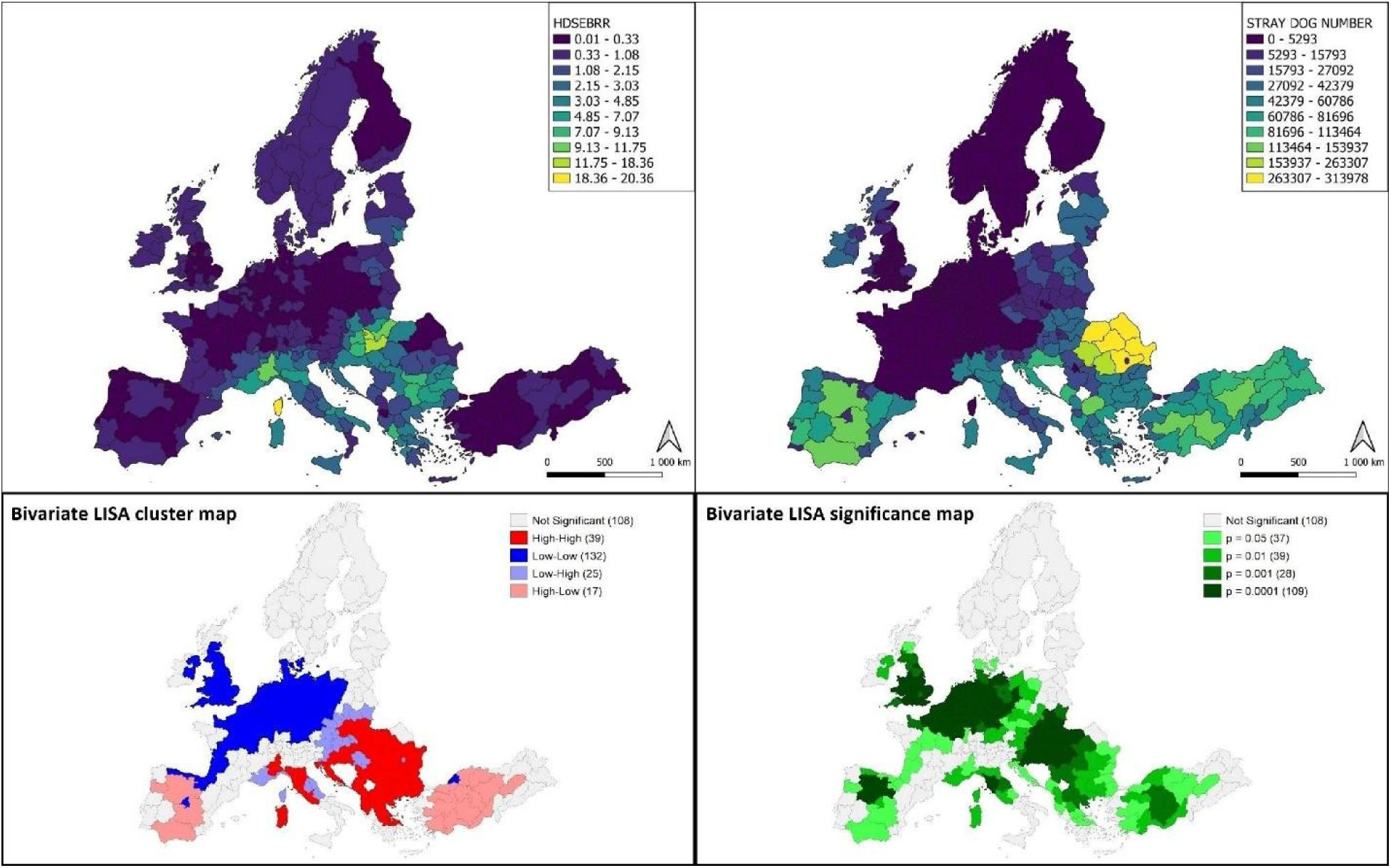
The spatial correlation between the spatial relative risk of human dirofilariasis and the estimated number of stray dogs at the NUTS 2 level. (Note: Top left: HDSEBRR (spatial Empirical Bayes relative risk of human *Dirofilaria repens* infection). Top right: estimated number of stray dogs by region (STRAY DOG NUMBER) (derived from estimative data using area-weighted interpolation, original data: https://www.esdaw.eu/stray-animals-by-country.html). Bottom left: bivariate LISA cluster map (High-High: high number of stray dogs and high infection risk; Low-Low: low number of stray dogs and low infection risk; Low-High: low number of stray dogs, high infection risk; High-Low: high number of stray dogs, low infection risk; number of regions in parentheses). Bottom right: bivariate LISA significance map (number of regions per significance level in parentheses)).

The LISA cluster map shows a pattern similar to that of the deprivation analysis, but even more pronounced. The number of High-High clusters is higher (39 regions, compared to 28 regions in the deprivation analysis), and the Low-Low clusters are also more widespread (132 regions, compared to 110). This suggests that the number of stray dogs shows a stronger spatial association with the risk of human dirofilariasis than social deprivation alone does.

High-High clusters are concentrated in Southeast Europe—in Romania, Bulgaria, Greece, and other Balkan countries—where large numbers of stray dogs serve as a natural reservoir for the parasite. Stray dogs are the primary definitive hosts of *D. repens*, and as reservoirs of microfilariae, they serve as sources of infection for mosquito vectors. High-low clusters can be found in large parts of Spain and Türkiye, where the inappropriate, too hot and dry climate impedes the circulation of infection despite the dense stray dog populations.

## Discussion

Human dirofilariasis caused by *D. repens* exhibits an expanding range within the Palaearctic [1,47]. Although the clinical manifestations of the infection are predominantly mild, lesions affecting the eye, internal organs [69,70] or the central nervous system [71] may generate severe complications. Besides climatic factors, socioeconomic conditions are increasingly suspected as drivers of disease transmission [19,23], thereby altering the parasite’s geographic range. Consequently, the northward expansion is not uniform; certain European regions appear to have a high risk despite a dry continental climate with harsh winters. This is evidenced by case clusters observed in Kyiv and Dnipropetrovsk, Ukraine [72], Sofia, Bulgaria [18], and along the Austrian-Hungarian border [73]. Specifically, the Carpathian Basin has been proposed as a central corridor facilitating the successful range expansion of *D. repens*, given that the majority of cases in both Slovakia and Austria have been concentrated along their shared borders with Hungary, the central country of the Basin.

Based on the literature, we hypothesised that the Carpathian Basin plays a significant role in channelling *D. repens* northwards from the Balkans into the continental interior. Furthermore, our sub-hypotheses were that both climatic and socioeconomic factors facilitate this process. To test these hypotheses, published data on European cases [1], along with 145 Hungarian cases, were analysed using MaxEnt models to identify the potential ecological drivers of *D. repens* transmission across Europe. In addition, infection risks within specific European administrative units were correlated with their respective social deprivation indices and estimated stray dog populations.

The MaxEnt models revealed that the predicted probability of human *D. repens* infection within the Carpathian Basin is as high as that of the Adriatic and Black Sea coastal area (Figure 6). The primary drivers of human infection risk include summer warmth (MWMT), winter coldness (Tmin_wt), and summer precipitation (Prec_sm). While low winter temperature acts as an absolute limiting factor with a lower threshold of -2.1 °C, warmth and precipitation have optimal ranges (Figure 5). The summer thermal optimum, with values of 22.8-25.1 °C, aligns with the findings that hyperendemic regions in Mediterranean countries are located away from the hottest subregions. In Greece, for instance, the most heavily affected region [74] lies in the north, bordering Bulgaria’s inland region, which is also a known hotspot [18]. Similarly, in Italy, the highest number of human cases was reported in the northernmost regions of the Po Plain, while the southernmost regions appear to be less affected [1] (Figure 3).

These findings correspond to studies on mosquito heat tolerance. Investigations have demonstrated that culicids, as ectotherms, are insufficiently tolerant of high temperatures, and only their avoidance behaviour, seeking shelter, protects them from overheating during the hottest hours [75]. Mosquitoes are most vulnerable in xeric biomes, particularly within the subtropical climatic zone [76]. A decreasing suitability of the southernmost Mediterranean regions for mosquito vector species has also been observed in malaria. In parallel with the northward expansion of Anopheles occurrence, a decline is expected in the south. Consequently, a warming climate might induce a northward shift of the malaria epidemic belt over central-northern Europe [14].

The mosquito vectors of *D. repens* rely on wet habitats that offer optimal conditions for oviposition [5,77,78]. Furthermore, as ectotherms, they require environmental temperatures that align with their thermal tolerance [75,76]. Consequently, surface waters and surrounding wet habitats with shady forest vegetation are expected to provide ideal survival and breeding environments for potential vector species. However, in this study, the surface water variable was removed from the modelling process due to its low permutation importance.

Literature on culicids has revealed varied ecological requirements among specific potential vector species. Floodwater mosquitoes, like *Ae. vexans*, *Ae. caspius*, and *Ae. sticticus* [4,5,78] are closely linked to floodplain ecosystems, *Culex pipiens* biotypes depend on synanthropic environments [9,77,79], while *Anopheles* spp. prefer warm surface waters in the vicinity of stables and barns, which are primarily found in Mediterranean agroecosystems [13,15].

Furthermore, the most river-dependent floodwater mosquitoes possess the highest capacity for long-distance flight from their breeding sites when seeking for blood meals [3,80], enabling them to expand their range even into arid habitats [7]. The gonotrophic cycle of culicids lasts for 3-7 days, depending on ambient temperature [81,82]. This period comprises blood feeding, digestion, oogenesis, and oviposition. Only oviposition relies strictly on wet habitats, while feeding is closely linked to host availability. Therefore, mosquitoes capable of long-distance flight, such as robust *Aedes* species [3,81], contribute to disease risk even far from extended surface waters.

On the other hand, synanthropic culicids can utilise wet microhabitats that entrap water, such as rainwater containers, dumped plastic and porcelain debris, and discarded tyres. Consequently, they do not need to leave host-dense synanthropic environments to reproduce [83–85]. The ecological plasticity of certain species, combined with the wide variety of vectors [1], explains why our analyses revealed that the vicinity of surface waters is a less relevant driver of human *D. repens* infection. However, the visualisation of case distribution in the Carpathian Basin, Hungary (Figure 2) suggests that large river valleys exert an effect, albeit not statistically significant, on the occurrence of human cases.

Regarding precipitation, a summer optimum between 28.6 and 231 mm was calculated. The lower limit is consistent with previous findings indicating that mosquitoes can poorly tolerate summer heat in xeric shrublands of the subtropical climate zone [76], a description that fits the southernmost parts of Greece and Spain, as demonstrated by the map in Figure 6. The impact of the upper limit is detectable in northwestern Europe under an Atlantic climate. High precipitation throughout the year may reduce immature culicid populations by flushing out breeding sites and diminishing the bacterial food sources of mosquito larvae [86].

Our study found a positive correlation between the risk of human *D. repens* infection and social deprivation in the southern part of Central and Eastern Europe (CEE), primarily the Balkan countries and the Carpathian Basin (Figure 7). A very similar pattern became apparent regarding the relationship between infection risk and stray dog population density (Figure 8). In the western part of the continent, such a definitive correlation between these factors and disease risk was not detected. However, it is critical to note that the calculation of the DEPRIVATION variable was based on the average social status within each specific country. Therefore, “socioeconomic vulnerability” does not represent the same standard of living in a Western European country as it does in a former Soviet satellite state. The post-communist bloc countries have been unable to overcome their structural lag over the past decades. Especially in Romania, Bulgaria, and Hungary, general economic growth has been accompanied by rising inequality [87].

The high abundance of stray dogs in CEE countries may be a consequence of socioeconomic factors, as the livelihood issues of owners have been identified as the primary cause of dog abandonment. Nevertheless, developed Mediterranean countries, such as Italy and Spain, also harbour significant populations of stray dogs [88]. Financial difficulties of dog owners [89] and the overwhelmed capacities of animal shelters [90] (Natoli et al., 2019) are again suspected to underpin this phenomenon. Since several Western European welfare states consider themselves free of stray dogs, regionally varying correlations between the parasite’s risk and the number of unowned dogs in those areas is not surprising (Figure 8). The role of stray dogs as reservoirs for canine dirofilariasis has already been raised [23,91,92]. Although the European maps of stray dog populations (Figure 8) and social deprivation (Figure 7) display dissimilar regional patterns, a large-scale global survey across 14 countries revealed a robust relationship between social well-being and animal welfare, independent of the respondents’ cultural background [93].

However, poverty does not solely contribute to dog abandonment. Studies investigating the association between waste management and social vulnerability have highlighted a strong bidirectional link. Marginalised societal strata often contribute to substandard waste management due to lower educational attainment and lacking financial resources to invest in sanitation [94–96]. The disadvantageous social effects of landfill siting can be observed even in higher-income countries. Although decision-makers frequently select already deprived areas for landfill siting, the establishment of a new landfill further contributes to regional backwardness [97]. In CEE countries, waste management policies have exacerbated social inequalities, as landfills operating with outdated technologies are predominantly sited close to the residential areas of socioeconomically disadvantaged groups [98,99]. The association between unimproved waste management and diseases, particularly vector-borne diseases, is well documented [84,100].

Our study investigated the potential role of the Carpathian Basin as a facilitator for the range expansion of the *D. repens* toward the interior of the European continent. The MaxEnt analyses highlighted a high probability of human *D. repens* infection within the Basin (Figure 6). Our findings align with the long-standing theory of the “Illyric-Dacian pincer”, which was established in the early 21th century. Based on the investigation of endemic invertebrates of the Balkans and the Carpathian Basin, researchers proved that the two ecoregions are interconnected, allowing for transition into one another. Consequently, endemic species of the Balkans unimpededly expand into the Basin [30].

The Basin encompasses the middle reaches of two major rivers, the Danube and the Tisza. Despite the MaxEnt models indicating a limited permutation relevance of surface waters, the visualisation of infection probability in Figure 6 delineates a high-risk corridor along the Danube River, extending from its Black Sea delta to its origin in Germany. A remarkably similar pattern is observable along the Tisza River, which appears to extend the high-risk zone to the eastern slopes of the Ukrainian Carpathians. Interestingly, the regional case distribution presented in Figure 2 suggests an elevated risk along the Tisza River, while the MaxEnt analysis utilising a larger sample size highlights that the probability of *D. repens* infection is higher in the western part of the Carpathian Basin. This finding aligns with the conclusions of a previous study on canine dirofilariasis. Those researchers found that the occurrence of *D. repens* is more probable in the Danube River catchment, with a definitive cluster in the southwestern part of the country [101].

Between France and the Black Sea, a contiguous mountain range isolates the continental interior from the Mediterranean Basin. The only ecological corridor is the Danube Valley that breaks through the narrow gap between the Alps and the Carpathians forming a migration route for the biota comprising plants, animals, and even prehistoric humans [26,27]. The strong isolation effect of mountain ranges and the corridor role of the Danube are observable in the Figure 4 map generated with MaxEnt analysis allowing us to extend the Danube corridor theory to the expansion of subtropical vector-borne parasites [26].

Beyond ecological suitability, the Carpathian Basin constitutes the central part of the CEE, where the economic development driven by European Union (EU) membership has failed to eradicate social inequality and the segregation of low-income communities [87]. Consequently, large populations continue to live on the periphery of society. These marginalised groups, characterised by lower educational attainment and limited financial resources, are unable to invest in sanitation and healthcare [38,39], and particularly not in the health of their dogs [93]. In Hungary, the central country of the Carpathian Basin, the current administration has taken office with the stated goal of eradicating poverty and developing education and healthcare [102,103]. However, the preceding one and a half decades under the hybrid electoral authoritarian Orbán regime have significantly widened the educational attainment gap between societal strata, resulting in the long-term conservation of inequality [38], which may facilitate the silent spread of infectious diseases.

Most cases of human dirofilariasis are detected accidentally, and diagnosis relies primarily on morphological identification of surgically removed parasites [69]. Cases presenting with mild symptoms are usually neglected for months, even by the patients themselves [104]. Similarly, most canine cases also display barely noticeable signs [105]. These inherent features of the parasitic disease pose a significant challenge to surveillance attempts. Furthermore, the cryptic reservoir, comprising stray dogs and dogs owned by marginalised social groups, facilitates the long-term maintenance of *D. repens*, thereby increasing the risks from the ecological suitability of the Carpathian Basin.

This study is subject to certain limitations. The baseline dataset included human cases diagnosed across Europe. Considering the difficulties in diagnosis [69,104,106], obtaining accurate statistics on human *D. repens* infection remains elusive. Consequently, the data used in this study very likely underestimate the true epidemiological situation. An analysis of the map in Figure 3 reveals that densely populated cities appear to be more heavily affected than rural regions. This highlights an urban bias: in densely populated urban areas, rare diseases are more frequently detected, thereby creating an analytical artefact. In this instance, MaxEnt seemingly mitigated the over-representation of urban patients, as the predicted probability map of human cases in Figure 6 does not display an elevated risk specifically in cities. However, the relevance of surface waters to *D. repens* epidemiology was evaluated as low, despite river valleys being conspicuously outlined on the map as higher-risk areas than their surroundings. This phenomenon might reflect the under-representation of rural patients, whose infection data could have clarified the true risk associated with wet habitats. Due to the lack of patient-specific clinical and epidemiological data, individual risk factors, such as educational attainment, poverty status, immunocompromising factors, could not be identified. In a future case-control study, such data will be indispensable for a better understanding of the transmission dynamics of this parasite.

## Conclusions

This study aimed to determine the most significant ecological and socioeconomic factors driving the epidemiology of human dirofilariasis caused by *D. repens* in Europe, with special focus on the role of the Carpathian Basin as a potential corridor facilitating the northward range expansion of the parasite into the continental interior. Employing MaxEnt analysis, the mean warmest month temperature (MWMT) was found to contribute primarily to the predicted probability of human infection. To a lesser extent, minimum winter temperature (Tmin_wt) and summer precipitation (Prec_sm) also affected the epidemiology of *D. repens* in Europe. Beyond climatic factors, social deprivation and stray dog population density influenced infection risk, though these trends varied spatially. The strongest positive correlation between disadvantageous social factors and increased health risk was detected in the eastern Balkan countries and within the Carpathian Basin. Furthermore, this continental region exhibits an ecological suitability for the *D. repens* circulation that is very similar to the Adriatic and the Black Sea coastal areas. Consequently, by integrating the ecological and socioeconomical findings of our study, the Carpathian Basin emerges as a high-risk area for *D. repens* transmission northwards.

## List of abbreviations

AICc: corrected Akaike Information Criterion
AUC: Area Under the (Receiver Operating Characteristic) Curve
AUC SD: AUC Standard Deviation
BiLISA: Bivariate Local Indicators of Spatial Association
CBI: Continuous Boyce Index
CEE: Central and Eastern Europe
CMD: Hargreaves climatic moisture deficit
ENM: Ecological Niche Model
EU: European Union
FC: feature class
H: hinge
HDSEBRR: relative risk of human *Dirofilaria repens* estimated by spatial Empirical Bayes smoothing
L: linear
LH: linear-hinge
LQ: linear-quadratic
LQH: linear-quadratic-hinge
MCMT: mean coldest month temperature (°C)
MSP: mean summer (May to Sept.) precipitation (mm)
MWMT: mean warmest month temperature (°C)
ncoef: number of coefficients
NLR: National Reference Laboratory of Medical Parasitology (Department of Bacteriology, Mycology and Parasitology, National Centre for Public Health and Pharmacy, Hungary)
NUTS 2: second level of the Nomenclature of Territorial Units for Statistics
PC: per cent contribution
PI: permutation importance
Prec_at: autumn precipitation (mm)
Prec_sm: summer precipitation (mm)
Prec_sp: spring precipitation (mm)
Q: quadratic
RM: regularisation multiplier/ regularisation factor
ROC: Receiver Operating Characteristic
SHM: summer heat-moisture index ((MWMT)/(MSP/1000))
SW_prop: surface water proportion (the summarised proportion of water courses (CLC2018 class: 511) and water bodies (CLC2018 class 512))
Tmax_at: autumn mean maximum temperature (°C)
Tmax_sm: summer mean maximum temperature (°C)
Tmax_sp: spring mean maximum temperature (°C)
Tmin_wt: winter mean minimum temperature (°C)
TSS: True Skill Statistic
VIF: variance-inflation factor
w.AIC: Akaike Information Criterion weight

## Declarations

### Ethics approval and consent to participate

Not applicable.

### Consent for publication

Not applicable.

### Availability of data and materials

The datasets supporting the conclusions of this article are available in the Zenodo repository, https://zenodo.org/records/21054890.

### Authors’ contributions

Conceptualisation: IK, TS. Investigation, data curation, writing - original draft: EN, IZ, ÁC. Data curation, visualisation, data management: TT, GN. Writing - review & editing: ST, ÁC, GN.

## Acknowledgements

During the preparation of this manuscript, the authors used Google Gemini 1.5 Pro (version 3.5) and Grammarly for the purposes of correcting grammatical errors and improving fluency. The authors have reviewed and edited the output and take full responsibility for the content of this publication.

## References

1. Hattendorf C, Lühken R. Vectors, host range, and spatial distribution of *Dirofilaria immitis* and *D. repens* in Europe: a systematic review. Infect Dis Poverty. 2025;14(1):58. 10.1186/s40249-025-01328-2.

2. Simón F, Siles-Lucas M, Morchón R, González-Miguel J, Mellado I, Carretón E, Montoya-Alonso JA. Human and animal dirofilariasis: the emergence of a zoonotic mosaic. Clin Microbiol Rev. 2012;25(3):507–44. 10.1128/cmr.00012-12.

3. Vaux AGC, Watts D, Findlay-Wilson S, Johnston C, Dallimore T, Drage P, Medlock JM. Identification, surveillance and management of *Aedes vexans* in a flooded river valley in Nottinghamshire, United Kingdom. J Eur Mosquito Control Assoc. 2021;39(1):15–25. 10.52004/JEMCA2021.0001.

4. Calzolari M, Mosca A, Montarsi F, Grisendi A, Scremin M, Roberto P, Tessarolo C, Defilippo F, Gobbo F, Casalone C, Lelli D, Albieri A. Distribution and abundance of *Aedes caspius* (Pallas, 1771) and *Aedes vexans* (Meigen, 1830) in the Po Plain (northern Italy). Parasites Vectors. 2024;17(1):452. 10.1186/s13071-024-06527-8.

5. Schäfer ML, Lundström JO. Different responses of two floodwater mosquito species, *Aedes vexans* and *Ochlerotatus sticticus* (Diptera: Culicidae), to larval habitat drying. J Vector Ecol. 2006;31(1):123–8. 10.3376/1081-1710(2006)31[123:DROTFM]2.0.CO;2.

6. Novak RJ. Oviposition sites of *Aedes vexans* (Diptera: Culicidae): wet-prairie habitats. Can Entomol. 1981;113(1):57–64. 10.4039/Ent11357-1.

7. Outammassine A, Zouhair S, Loqman S. Global potential distribution of three underappreciated arboviruses vectors (*Aedes japonicus*, Aedes vexans and Aedes vittatus) under current and future climate conditions. Transbound Emerg Dis. 2022;69(4):e1160–e1171. 10.1111/tbed.14404.

8. Becker N, Jöst A, Weitzel T. The *Culex pipiens* complex in Europe. J Am Mosquito Control Assoc. 2012;28(4s):53–67. 10.2987/8756-971X-28.4s.53.

9. Haba Y, McBride L. Origin and status of *Culex pipiens* mosquito ecotypes. Curr Biol. 2022;32(5):R237–R246. 10.1016/j.cub.2022.01.062.

10. Farajollahi A, Fonseca DM, Kramer LD, Kilpatrick AM. “Bird biting” mosquitoes and human disease: a review of the role of *Culex pipiens* complex mosquitoes in epidemiology. Infect Genet Evol. 2011;11(7):1577–85. 10.1016/j.meegid.2011.08.013.

11. Kim S, Trocke S, Sim C. Comparative studies of stenogamous behaviour in the mosquito *Culex pipiens* complex. Med Vet Entomol. 2018;32(4):427–35. 10.1111/mve.12309

12. Vinogradova EB. *Culex pipiens pipiens* mosquitoes: taxonomy, distribution, ecology, physiology, genetics, applied importance and control (Mosquitoes Monographs, No. 2). Sofia: Pensoft Publishers; 2000.

13. Gilioli G, Defilippo F, Simonetto A, Heinzl A, Migliorati M, Calzolari M, Canziani S, Lelli D, Lavazza A. Characterization of environmental drivers influencing the abundance of *Anopheles maculipennis* complex in Northern Italy. Parasites Vectors. 2024;17(1):109. 10.1186/s13071-024-06208-6

14. Hertig E. Distribution of *Anopheles* vectors and potential malaria transmission stability in Europe and the Mediterranean area under future climate change. Parasites Vectors. 2019;12(1):18. 10.1186/s13071-018-3278-6

15. Bertola M, Mazzucato M, Pombi M, Montarsi F. Updated occurrence and bionomics of potential malaria vectors in Europe: a systematic review (2000–2021). Parasites Vectors. 2022;15(1):88. 10.1186/s13071-022-05204-y

16. Lühken R, Becker N, Dyczko D, Sauer FG, Kliemke K, Schmidt-Chanasit J, Rydzanicz K. First record of *Anopheles* (*Anopheles*) *hyrcanus* (Pallas 1771) (Diptera: Culicidae) in Poland. Parasites Vectors. 2023;16(1):345. 10.1186/s13071-023-05974-z

17. Ciucă L, Genchi M, Kramer L, Mangia C, Miron LD, Del Prete L, Cringoli G, Rinaldi L. Heat treatment of serum samples from stray dogs naturally exposed to *Dirofilaria immitis* and *Dirofilaria repens* in Romania. Vet Parasitol. 2016;225:81–5. 10.1016/j.vetpar.2016.05.032

18. Harizanov RN, Jordanova DP, Bikov IS. Some aspects of the epidemiology, clinical manifestations, and diagnosis of human dirofilariasis caused by *Dirofilaria repens*. Parasitol Res. 2014;113(4):1571–9. 10.1007/s00436-014-3802-3

19. Otranto D, Dantas-Torres F, Mihalca AD, Traub RJ, Lappin M, Baneth G. Zoonotic parasites of sheltered and stray dogs in the era of the global economic and political crisis. Trends Parasitol. 2017;33(10):813–25. 10.1016/j.pt.2017.05.013

20. de Jesús Crespo R, Rogers RE. Habitat segregation patterns of container breeding mosquitos: the role of urban heat islands, vegetation cover, and income disparity in cemeteries of New Orleans. Int J Environ Res Public Health. 2022;19(1):245. 10.3390/ijerph19010245

21. Bowman DD, Liu Y, McMahan CS, Nordone SK, Yabsley MJ, Lund RB. Forecasting United States heartworm *Dirofilaria immitis* prevalence in dogs. Parasites Vectors. 2016;9(1):540. 10.1186/s13071-016-1804-y.

22. Chocobar MLE, Schmidt EMDS, Mendes ÂJF, Johnson PCD, Weir W, Panarese R. Microgeographical variation in *Dirofilaria immitis* prevalence in dogs in suburban and urban areas of Rio de Janeiro, Brazil. Vet Sci. 2025;12(1):3. 10.3390/vetsci12010003.

23. Morchón R, Balmori-de la Puente A, Infante González-Mohino E, Esteves-Guimarães J, Busquets P, Fontes-Sousa AP, Carretón E, Montoya-Alonso JA. Deciphering the socio-environmental factors associated with realized heartworm transmission risk in dogs from Portugal and Spain. Front Vet Sci. 2026;13:1812406. 10.3389/fvets.2026.1812406.

24. Gaudenyi T, Mihajlović M. The Carpathian Basin: denomination and delineation. Eur J Environ Earth Sci. 2022;3(2):1–6.

25. Jánosi IM, Bíró T, Lakatos BO, Gallas JA, Szöllosi-Nagy A. Changing water cycle under a warming climate: tendencies in the Carpathian Basin. Climate. 2023;11(6):118. 10.3390/cli11060118.

26. Chu W. The Danube corridor hypothesis and the Carpathian Basin: geological, environmental and archaeological approaches to characterizing Aurignacian dynamics. J World Prehist. 2018;31(2):117–78. 10.1007/s10963-018-9115-1.

27. Saag L, Staniuk R. Historical human migrations: from the steppe to the basin. Curr Biol. 2022;32(13):R738–R741. 10.1016/j.cub.2022.05.058.

28. Gömöry D, Zhelev P, Brus R. The Balkans: a genetic hotspot but not a universal colonization source for trees. Plant Syst Evol. 2020;306(1):5. 10.1007/s00606-020-01647-x.

29. Španiel S, Rešetnik I. Plant phylogeography of the Balkan Peninsula: spatiotemporal patterns and processes. Plant Syst Evol. 2022;308(5):38. 10.1007/s00606-022-01831-1.

30. Mahunka S, Murányi D, Kontschán J. The role of the Balkan Peninsula in the origin and genesis of the soil fauna of the Carpathian Basin: history, aims and results. Opusc Zool. 2013;44(Suppl 1):5–10.

31. Umhang G, Bastid V, Avcioglu H, Bagrade G, Bujanić M, Čabrilo OB, Casulli A, Dorny P, van der Giessen J, Guven E, Harna J, Karamon J, Kharchenko V, Knapp J, Kolarova L, Konyaev S, Laurimaa L, Losch S, Miljević M, Miterpakova M, Moks E, Romig T, Saarma U, Snabel V, Sreter T, Valdmann H, Boué F. Unravelling the genetic diversity and relatedness of *Echinococcus multilocularis* isolates in Eurasia using the EmsB microsatellite nuclear marker. Infect Genet Evol. 2021;92:104863. 10.1016/j.meegid.2021.104863.

32. Kiss K, Bálint M, Gémes A, Marcsik A, Dávid Á, Évinger S, Hajdu T, Szeniczey T. More than one millennium (2nd-16th century CE) of the White Plague in the Carpathian Basin–New cases, expanding knowledge. Tuberculosis. 2023;143:102387. 10.1016/j.tube.2023.102387.

33. Nagy E, Nagy RR, Miklós M, Szekeres S, Abdalrahman BM, Földvári G, Hornok S, Nagy G. Eye to eye with *Thelazia*-infected canids in Central European forests. Curr Res Parasitol Vector Borne Dis. 2026;100353. 10.1016/j.crpvbd.2026.100353.

34. Rew LJ, McDougall KL, Alexander JM, Daehler CC, Essl F, Haider S, Kueffer C, Lenoir J, Milbau A, Nuñez MA, Pauchard A, Rabitsch W. Moving up and over: redistribution of plants in alpine, Arctic, and Antarctic ecosystems under global change. Arctic Antarctic Alpine Res. 2020;52(1):651–65. 10.1080/15230430.2020.1845919.

35. Ilona J, Bartók B, Dumitrescu A, Cheval S, Gandhi A, Tordai ÁV, Weidinger T. Using long-term historical meteorological data for climate change analysis in the Carpathian region. Atmosphere. 2022;13(11):1751. 10.3390/atmos13111751.

36. Skarbit N, Unger J, Gál TM. Projected values of thermal and precipitation climate indices for the broader Carpathian region based on EURO-CORDEX simulations. Hung Geogr Bull. 2022;71(4):325–47. 10.15201/hungeobull.71.4.2.

37. Didovets I, Krysanova V, Bürger G, Snizhko S, Balabukh V, Bronstert A. Climate change impact on regional floods in the Carpathian region. J Hydrol Reg Stud. 2019;22:100590. 10.1016/j.ejrh.2019.01.002.

38. Czibere I, Balogh K, Kovách I, Nemes-Zámbó G. Exclusionary Mechanisms of Social Policy Redistribution in Hungary. Soc Policy Soc. 2026;25(1):92–107. 10.1017/S1474746424000149.

39. Szombati K. The consolidation of authoritarian rule in rural Hungary: Workfare and the shift from punitive populist to illiberal paternalist poverty governance. Eur Asia Stud. 2021;73(9):1703–25. 10.1080/09668136.2021.1990861.

40. Kovács JM, Trencsényi B. Conclusion: Hungary—Brave and new? Dissecting a realistic dystopia. In: Kovács JM, Trencsényi B, editors. Brave New Hungary: Mapping the “System of National Cooperation”. Lanham: Lexington Books; 2020. p. 379–432.

41. Neumann E. How churches make education policy: the churchification of Hungarian education and the social question under religious populism. Religion State Soc. 2025;53(2):97–116. 10.1080/09637494.2024.2399452.

42. Phillips SJ, Anderson RP, Schapire RE. Maximum entropy modelling of species geographic distributions. Ecol Modell. 2006;190:231–59.

43. Carvalho BM, Rangel EF, Vale MM. Evaluation of the impacts of climate change on disease vectors through ecological niche modelling. Bull Entomol Res. 2017;107(4):419–30. 10.1017/S0007485316001097

44. Escobar LE. Ecological Niche Modeling: An Introduction for Veterinarians and Epidemiologists. Front Vet Sci. 2020;7:519059. 10.3389/fvets.2020.519059

45. Lawrence TJ, Takenaka BP, Garg A, Tao D, Deem SL, Fèvre EM, Gluecks I, Sagan V, Shacham E. A global examination of ecological niche modeling to predict emerging infectious diseases: a systematic review. Front Public Health. 2023;11:1244084. 10.3389/fpubh.2023.1244084

46. Araujo MB, Guisan A. Five (or so) challenges for species distribution modelling. J Biogeogr. 2006;33:1677–88.

47. Capelli G, Genchi C, Baneth G, Bourdeau P, Brianti E, Cardoso L, Danesi P, Fuehrer HP, Giannelli A, Ionică AM, Maia C, Modrý D, Montarsi F, Krücken J, Papadopoulos E, Petrić D, Pfeffer M, Savić S, Otranto D, Poppert S, Silaghi C. Recent advances on *Dirofilaria repens* in dogs and humans in Europe. Parasites Vectors. 2018;11(1):663. 10.1186/s13071-018-3205-x.

48. Kucsera I, Danka J, Fok É. A dirofilariosis közegészségügyi és állategészségügyi jelentősége Magyarországon: Legújabb ismeretek a dirofilariosisról (The public health and veterinary importance of dirofilariasis in Hungary: Latest knowledge on dirofilariasis). Mikrobiol Körlevél. 2021;21(2):142–58.

49. Sillero N, Barbosa AM. Common mistakes in ecological niche models. Int J Geogr Inf Sci. 2021;35(2):213–26. 10.1080/13658816.2020.1798968.

50. Sillero N, Arenas-Castro S, Enriquez-Urzelai U, Gomes Vale C, Sousa-Guedes D, Martínez-Freiría F, Real R, Barbosa MA. Want to model a species niche? A step- by-step guideline on correlative ecological niche modelling. Ecol Modell. 2021;456:109671. 10.1016/j.ecolmodel.2021.109671.

51. Naimi B, Hamm NA, Groen TA, Skidmore AK, Toxopeus AG. Where is positional uncertainty a problem for species distribution modelling. Ecography. 2014;37(2):191 –203. 10.1111/j.1600-0587.2013.00205.x

52. Hijmans RJ, Brown A, Barbosa M. terra: Spatial Data Analysis. R package version 1.9–34. 2026. 10.32614/CRAN.package.terra.

53. Kass JM, Muscarella R, Galante PJ, Bohl CL, Buitrago-Pinilla G, Boria RA, Soley-Guardia M, Anderson RP. ENMeval 2.0: Redesigned for customizable and reproducible modeling of species’ niches and distributions. Methods Ecol Evol. 2021;12(9):1602–8. 10.1111/2041-210X.13628

54. Cobos ME, Peterson AT, Barve N, Osorio-Olvera L. kuenm: an R package for detailed development of ecological niche models using Maxent. PeerJ. 2019;7:e6281. 10.7717/peerj.6281

55. Urbanek S. rJava: Low-Level R to Java Interface. R package version 1.0–18. 2026. 10.32614/CRAN.package.rJava.

56. Burnham KP, Anderson DR. Model Selection and Multimodel Inference: A Practical Information-Theoretic Approach. 2nd ed. New York: Springer; 2002.

57. Swets JA. Measuring the accuracy of diagnostic systems. Science. 1988;240(4857):1285–93.

58. Hirzel AH, Le Lay G, Helfer V, Randin C, Guisan A. Evaluating the ability of habitat suitability models to predict species presences. Ecol Modell. 2006;199(2):142 –52. 10.1016/j.ecolmodel.2006.05.017.

59. Radosavljevic A, Anderson RP. Making better Maxent models of species distributions: complexity, overfitting and evaluation. J Biogeogr. 2014;41(4):629–43.

60. Merow C, Smith MJ, Silander JA Jr. A practical guide to MaxEnt for modeling species’ distributions: what it does, and why inputs and settings matter. Ecography. 2013;36(10):1058–69.

61. Whitford AM, Shipley BR, McGuire JL. The influence of the number and distribution of background points in presence-background species distribution models. Ecol Modell. 2024;488:110604. 10.1016/j.ecolmodel.2023.110604.

62. Allouche O, Tsoar A, Kadmon R. Assessing the accuracy of species distribution models: prevalence, kappa and the true skill statistic (TSS). J Appl Ecol. 2006;43(6):1223–32.

63. Liu C, Newell G, White M. On the selection of thresholds for predicting species occurrence with presence-only data. Ecol Evol. 2016;6(2):337–48.

64. Austin MP. Spatial prediction of species distribution: an interface between ecological theory and statistical modelling. Ecol Modell. 2002;157(2–3):101–18. 10.1016/S0304-3800(02)00205-3.

65. Austin M. Species distribution models and ecological theory: A critical assessment and some possible new approaches. Ecol Modell. 2007;200(1–2):1–19. 10.1016/j.ecolmodel.2006.07.005.

66. Citores L, Ibaibarriaga L, Lee DJ, Brewer MJ, Santos M, Chust G. Modelling species presence–absence in the ecological niche theory framework using shape-constrained generalized additive models. Ecol Modell. 2020;418:108926. 10.1016/j.ecolmodel.2019.108926.

67. Anselin L, Syabri I, Kho Y. GeoDa: An Introduction to Spatial Data Analysis. Geogr Anal. 2006;38(1):5–22.

68. Anselin L. Exploring Spatial Data with GeoDa™: A Workbook. Urbana: Spatial Analysis Laboratory, Department of Geography, University of Illinois, Urbana-Champaign; 2005.

69. Miterpáková M, Antolová D, Ondriska F, Gál V. Human *Dirofilaria repens* infections diagnosed in Slovakia in the last 10 years (2007–2017). Wien Klin Wochenschr. 2017;129(17–18):634–41. 10.1007/s00508-017-1233-8.

70. Boldiš V, Ondriska F, Bošák V, Hajdúk O, Antolová D, Miterpáková M. Pseudo-tumor of the epididymis, a rare clinical presentation of human *Dirofilaria repens* infection: a report of autochthonous case of dirofilariasis in Southwestern Slovakia. Acta Parasitol. 2020;65(2):550–3. 10.2478/s11686-020-00170-w.

71. Poppert S, Hodapp M, Krueger A, Hegasy G, Niesen WD, Kern WV, Tannich E. *Dirofilaria repens* infection and concomitant meningoencephalitis. Emerg Infect Dis. 2009;15(11):1844–6. 10.3201/eid1511.090936.

72. Sa³amatin RV, Pavlikovska TM, Sagach OS, Nikolayenko SM, Kornyushin VV, Kharchenko VO, Masny A, Cielecka D, Konieczna-Sałamatin J, Conn D, Golab E. Human dirofilariasis due to *Dirofilaria repens* in Ukraine, an emergent zoonosis: epidemiological report of 1465 cases. Acta Parasitol. 2013;58(4):592–8. 10.2478/s11686-013-0187-x.

73. Geissler N, Ruff J, Walochnik J, Ludwig W, Auer H, Wiedermann U, Geissler W. Autochthonous human *Dirofilaria repens* infection in Austria. Acta Parasitol. 2022;67(2):1039–43. 10.1007/s11686-021-00506-0.

74. Dimzas D, Aindelis G, Tamvakis A, Chatzoudi S, Chlichlia K, Panopoulou M, Diakou A. *Dirofilaria immitis* and *Dirofilaria repens*: investigating the prevalence of zoonotic parasites in dogs and humans in a hyperenzootic area. Animals. 2024;14(17):2529. 10.3390/ani14172529.

75. Sunday JM, Bates A. E, Kearney MR, Colwell RK, Dulvy NK, Longino JT, Huey RB. Thermal-safety margins and the necessity of thermoregulatory behavior across latitude and elevation. Proc Natl Acad Sci U S A. 2014;111(15):5610–5. 10.1073/pnas.1316145111.

76. Couper LI, Nalukwago DU, Lyberger KP, Farner JE, Mordecai EA. How much warming can mosquito vectors tolerate?. Glob Change Biol. 2024;30(12):e17610. 10.1111/gcb.17610.

77. Marcolin L, Zardini A, Longo E, Caputo B, Poletti P, Di Marco M. Mapping the habitat suitability of *Culex pipiens* in Europe using ensemble bioclimatic modelling. J Biogeogr. 2025;52(10):e70019. 10.1111/jbi.70019.

78. Lindström A, Eklöf D, Lilja T. Different hatching rates of floodwater mosquitoes *Aedes sticticus*, *Aedes rossicus* and *Aedes cinereus* from different flooded environments. Insects. 2021;12(4):279. 10.3390/insects12040279.

79. Blom R, Krol L, Langezaal M, Schrama M, Trimbos KB, Wassenaar D, Koenraadt CJM. Blood-feeding patterns of *Culex pipiens* biotype *pipiens* and *pipiens/molestus* hybrids in relation to avian community composition in urban habitats. Parasites Vectors. 2024;17(1):95. 10.1186/s13071-024-06186-9.

80. Bogojević MS, Merdić E, Bogdanović T. The flight distances of floodwater mosquitoes (*Aedes vexans*, *Ochlerotatus sticticus* and *Ochlerotatus caspius*) in Osijek, Eastern Croatia. Biologia. 2011;66(4):678–83. 10.2478/s11756-011-0073-7.

81. Klowden MJ, Briegel H. Mosquito gonotrophic cycle and multiple feeding potential: contrasts between *Anopheles* and *Aedes* (Diptera: Culicidae). J Med Entomol. 1994;31(4):618–22. 10.1093/jmedent/31.4.618.

82. Al-Rashidi HS, Alghamdi KM, Al-Otaibi WM, Al-Solami HM, Mahyoub JA. Effects of blood meal sources on the biological characteristics of *Aedes aegypti* and *Culex pipiens* (Diptera: Culicidae). Saudi J Biol Sci. 2022;29(12):103448. 10.1016/j.sjbs.2022.103448.

83. Achaga JA, Vezzani D. A methodological proposal to estimate the total abundance of immature mosquitoes in discarded tyres: *Aedes aegypti* and *Culex pipiens* as study cases. Acta Trop. 2024;260:107474. 10.1016/j.actatropica.2024.107474.

84. Banerjee S, Aditya G, Saha GK. Household wastes as larval habitats of dengue vectors: comparison between urban and rural areas of Kolkata, India. PLoS One. 2015;10(10):e0138082. 10.1371/journal.pone.0138082.

85. Leisnham PT, LaDeau SL, Saunders ME, Villena OC. Condition-specific competitive effects of the invasive mosquito *Aedes albopictus* on the resident *Culex pipiens* among different urban container habitats may explain their coexistence in the field. Insects. 2021;12(11):993. 10.3390/insects12110993.

86. Valdez LD, Sibona GJ, Diaz LA, Contigiani MS, Condat CA. Effects of rainfall on *Culex* mosquito population dynamics. J Theor Biol. 2017;421:28–38. 10.1016/j.jtbi.2017.03.024.

87. Pop TM. The future of income inequality in the European Union: Do economic growth and poverty matter?. Port Econ J. 2026:1–25. 10.1007/s10258-026-00285-4.

88. Papavasili TH, Kontogeorgos A, Mavrommati A, Sossidou EN, Chatzitheodoridis F. Review of stray dog management: dog days in the European countries. Bulg J Vet Med. 2024;27(2):322–42. 10.15547/bjvm.2022-0035.

89. Belo VS, Werneck GL, da Silva ES, Barbosa DS, Struchiner CJ. Population Estimation Methods for Free-Ranging Dogs: A Systematic Review. PLoS One. 2015;10(12):e0144830. 10.1371/journal.pone.0144830.

90. Natoli E, Cariola G, Dall’Oglio G, Valsecchi P. Considerations of ethical aspects of control strategies of unowned free-roaming dog populations and the No-Kill policy in Italy. J Appl Anim Ethics Res. 2019;1(2):216–22. 10.1163/25889567-12340014.

91. Pękacz M, Slivinska K, Vyniarska A, Basałaj K, Kalinowska A, Wesołowska A, Laskowska A, Kysterna O, Klietsov A, Miterpáková M, Mihalca AD, Gawor J, Kharchenko V, Zawistowska-Deniziak A. Molecular investigation of *Dirofilaria repens*, *Dirofilaria immitis* and *Acanthocheilonema reconditum* in stray dogs and cats in Ukraine. BMC Vet Res. 2025;21(1):438. 10.1186/s12917-025-04867-w.

92. Dumitrache MO, D’Amico G, Voiniţchi E, Maximenco S, Mircean V, Ionică AM. An epidemiological survey of *Dirofilaria* spp. and *Acanthocheilonema* spp. in dogs from the Republic of Moldova. Parasites Vectors. 2021;14(1):390. 10.1186/s13071-021-04891-3

93. Sinclair M, Lee NY, Hötzel MJ, de Luna MCT, Sharma A, Idris M, Derkley T, Li C, Islam MA, Iyasere OS, Navarro G, Ahmed AA, Khruapradab C, Curry M, Burns GL, Marchant JN. International perceptions of animals and the importance of their welfare. Front Anim Sci. 2022;3:960379. 10.3389/fanim.2022.960379.

94. Adeoti A, Obidi B. Poverty and preference for improved solid waste management attributes in Delta-State, Nigeria. J Rural Econ Dev. 2010;19:15–33. 10.22004/ag.econ.206858.

95. Ma J, Hipel KW. Exploring social dimensions of municipal solid waste management around the globe–A systematic literature review. Waste Manage. 2016;56:3–12. 10.1016/j.wasman.2016.06.041.

96. Iacoboaea C, Luca O, Șercăianu M, Aldea M, Păunescu M, Popescu AL. Towards Sustainable Modes for Remote Monitoring in Waste Management: A Study of Marginalized Urban Areas in Romania. Sustainability. 2024;16(6):2400. 10.3390/su16062400.

97. Richardson EA, Shortt NK, Mitchell RJ. The mechanism behind environmental inequality in Scotland: which came first, the deprivation or the landfill?. Environ Plan A. 2010;42(1):223–40. 10.1068/a41376.

98. Dunajeva J, Kostka J. Racialized politics of garbage: waste management in urban Roma settlements in Eastern Europe. Ethnic Racial Stud. 2022;45(1):90–112. 10.1080/01419870.2020.1863442.

99. Škobla D, Filčák R. Life next to a landfill: urban marginality, environmental injustice and the Roma. Race Class. 2024;65(4):74–91. 10.1177/03063968231203488.

100. Degroote S, Zinszer K, Ridde V. Interventions for vector-borne diseases focused on housing and hygiene in urban areas: a scoping review. Infect Dis Poverty. 2018;7(1):96. 10.1186/s40249-018-0477-5.

101. Farkas R, Mag V, Gyurkovszky M, Takács N, Vörös K, Solymosi N. The current situation of canine dirofilariosis in Hungary. Parasitol Res. 2020;119(1):129–35. 10.1007/s00436-019-06478-5.

102. Howard S. Hungary: Why healthcare reform was a cornerstone of victory over Orbán. BMJ. 2026;393:s749. 10.1136/bmj.s749.

103. Benedek I. Polarizing transition? Opposition strategies and the rise of Péter Magyar and the Respect and Freedom Party (TISZA) in Hungary. Comp Eur Polit. 2026;24(1):24. 10.1057/s41295-026-00460-z.

104. Suzuki J, Kobayashi S, Okata U, Matsuzaki H, Mori M, Chen KR, Iwata S. Molecular analysis of *Dirofilaria repens* removed from a subcutaneous nodule in a Japanese woman after a tour to Europe. Parasite. 2015;22:2. 10.1051/parasite/2015002.

105. Paździor-Czapula K, Otrocka-Domagała I, Myrdek P, Mikiewicz M, Gesek M. *Dirofilaria repens*—an etiological factor or an incidental finding in cytologic and histopathologic biopsies from dogs. Vet Clin Pathol. 2018;47(2):307–11. 10.1111/vcp.12597.

106. Riebenbauer K, Weber PB, Walochnik J, Karlhofer F, Winkler S, Dorfer S, Auer H, Valencak J, Laimer M, Handisurya A. Human dirofilariosis in Austria: the past, the present, the future. Parasites Vectors. 2021;14(1):227. 10.1186/s13071-021-04696-4.

